# Distinct and shared neural resources between processing of dynamic physical objects and spatial working memory

**DOI:** 10.64898/2026.07.10.737787

**Authors:** Samuel Maione, Shari Liu

## Abstract

In human brains, portions of the frontoparietal cortex respond when people reason about the physical world (e.g. track objects, make predictions), and when people engage in a variety of demanding tasks (e.g. working memory, motor inhibition). Here, we use functional neuroimaging to address an open question about whether the same neural resources support both mental functions. Twenty-eight human adults (*M*_age_ = 26.5y; 17 female; 26 right handed) were scanned using functional magnetic resonance imaging (fMRI) while they (i) tracked the dynamics of physical objects (vs social agents), and (ii) performed a hard (vs easy) spatial working memory task. For each participant, we identified functional regions of interest (fROIs) that were maximally engaged by each task (physical > social; hard > easy), and studied their engagement in the held out task. We report three pieces of evidence that physical > social fROIs are recruited during the spatial working memory task. First, fROIs maximally engaged during physical (vs social) processing responded just as strongly during hard (vs easy) spatial working memory. Second, people with strong hard > easy responses in one set of regions also tended to have strong physical > social responses in the other set. Third, people with strong hard > easy responses in physical > social fROIs, but not in hard > easy fROIs, performed more strongly at spatial working memory in the scanner. These findings suggest that portions of the frontoparietal cortex that preferentially respond to physical (vs social) stimuli are involved in functions beyond physical reasoning: either spatial processing specifically, or attentionally demanding tasks in general.

## 1. Introduction

Every day, we interact with objects and reason about them in sophisticated ways. We stack, throw and catch objects, appreciate their properties like mass and material, and track and make predictions about them even when they move out of view. An open question is whether these capacities rely on specific mental resources–those particular to physical reasoning, or general mental resources–those applicable to tasks from many domains. One fruitful approach for addressing this question has been to study the distinct vs shared neural mechanisms between physical reasoning and domain-general thought (Fischer et al., 2016; Kean et al., 2025; Osiurak et al., 2025). Here, we use this approach to (1) study the neural substrates for domain-general attentional demand and those for processing of physical events, (2) relate neural responses from one set of regions to the other, and (3) relate neural responses in these regions to behavioral performance within the same domain.

Portions of the frontal and parietal cortex respond when people perceive and reason about the dynamics of the physical world (Fischer et al., 2016; Kean et al., 2025; Liu et al., 2024; Osiurak et al., 2025). These regions respond when people make physical predictions (Fischer et al., 2016; Kean et al., 2025; Zbären et al., 2023), watch movies or read stories about physical interactions (Hauptman & Bedny, 2024; Jack et al., 2013; Karakose-Akbiyik et al., 2023; Liu et al., 2024; Martin & Weisberg, 2003), and reason about the causal relations between objects (Osiurak et al., 2025; Wende et al., 2013). Regions of the frontoparietal cortex also contain information about abstract variables that are uniquely relevant for intuitive physics, including object mass (Schwettmann et al., 2018) and relations between objects like stability, containment, and contact (Pramod et al., 2022; Pramod, Mieczkowski, et al., 2025). Furthermore, these regions respond preferentially to surprising physical events (and not surprising actions; Liu et al., 2024; Parris et al., 2009; Plikat et al., 2025), and contain information about predicted future physical states (Pramod, Mieczkowski, et al., 2025), suggesting their role in encoding both predictions and prediction errors for physical events in particular. Frontoparietal cortex in infants responds to mechanical interaction stimuli and spatiotemporal changes in object motion (Wilcox & Biondi, 2015), suggesting their role in dynamic object processing early in development.

What are the functions of these regions that are preferentially engaged during physical reasoning? One possibility is that they are involved in stimulus tracking, maintaining information in working memory, selection across multiple actions or representations, and other functions that are relevant for thinking in many domains. In support of this view, demanding tasks involving motor inhibition, arithmetic, working memory, and other forms of cognitive control evoke activity in the frontoparietal cortex, in a set of regions sometimes referred to as the multiple demand network (Duncan, 2010; Fedorenko et al., 2013; Jackson & Woolgar, 2018; Thompson-Schill et al., 2005). For example, when Osiurak et al (2025) scanned adults on a variety of tasks involving mechanical reasoning, as well as Raven’s Progressive Matrices (a domain-general fluid intelligence task), they report group-level overlap in responses in several regions in frontoparietal cortex across the two tasks. More sensitive analyses in individual participants confirm spatial overlap between neural responses for physical inference tasks and working memory tasks (Fischer et al., 2016; Kean et al., 2024). Another suggestive piece of evidence comes from the cognitive profile of Williams syndrome–a genetic condition in which patients experience impaired physical reasoning (Kamps et al., 2017), spatial processing (Landau et al., 2006; O’Hearn et al., 2005), and working memory (Menghini et al., 2010), and intact performance in other abstract reasoning tasks, including tasks involving theory of mind (Kamps et al., 2017; Karmiloff-Smith et al., 1995).

On the other hand, capacities for physical reasoning are dissociable from other capacities in behavior. Studies of individual differences reveal that variation in performance in intuitive physics (e.g. judging the stability of stack of objects, predicting the trajectory of an object under occlusion, inferring the relative mass of two interacting objects) is separable from variation in performance on mental rotation, spatial and verbal working memory (Mitko et al., 2024; Mitko & Fischer, 2020). The neural resources devoted to physical reasoning and attentional demand are also partially dissociable, even in group analyses. For example, Osiurak et al. (2025) reported overlap in responses between physical reasoning tasks and a Raven’s matrices in group-level analyses in the frontal and parietal cortex, but they also found one region in the left inferior parietal lobule that responded to physical tasks, ranging from physical prediction to tool use (see also Fischer & Mahon, 2021; Goldenberg & Spatt, 2009; Osiurak et al., 2009; Seifert et al., 2025), and not Raven’s matrices. In other studies, the amplitude and patterns of neural responses evoked by multiple working memory tasks and intuitive physics tasks are more similar to each other (within task) than between any given working memory and physics task (across tasks) (Fischer et al., 2016; Kean et al., 2025), suggesting a partial separation (if not outright dissociation) between these two functions. The voxels in frontoparietal cortex most responsive during physical reasoning vs working memory were also more correlated to themselves than each other at rest (Pramod, Hutchinson, et al., 2025), further suggesting their distinctive functions. Yet, in the absence of how the neural substrates for physical processing and endogenous attention relate to the behavior associated with these two functions (cf. Assem et al., 2020), open questions remain about the nature of these overlapping vs distinct neural responses.

### 1.1 Present work

In this paper, we use a subject-specific functional regions of interest (fROI) approach (Fedorenko et al., 2010) to study whether voxels in the frontal and parietal cortex that are preferentially engaged by tracking and making predictions about physical (vs social) entities are also preferentially engaged in hard (vs easy) spatial working memory, and vice versa. First, we aimed to replicate prior findings that partially distinct neural populations in the frontoparietal cortex are preferentially involved in physical reasoning vs general attentional demand (Kean et al., 2025; Osiurak et al., 2025; Pramod, Hutchinson, et al., 2025).

We then take two novel approaches to further study the relationship between these two functions in the frontoparietal cortex. First, we ask whether individual differences in neural responses during physical reasoning can predict individual differences in neural responses during spatial working memory. Second, we ask whether either of these individual differences predict behavioral performance in the same tasks.

To preview our results, we report several pieces of evidence that physical reasoning and spatial attentional demand are subserved by partially distinct neural substrates. At the same time, we found evidence that the neural substrates underlying physical (vs social) processing are involved in spatial working memory, but not vice versa. First, we replicated prior findings that fROIs maximally engaged by physical reasoning and spatial working memory overlap. Second, we found an asymmetry in the functions of these fROIs. fROIs maximally engaged by spatial attentional demand do not show strong preferential responses for physical vs social content, suggesting that they are not preferentially engaged for physical reasoning. By contrast, fROIs maximally engaged by physical reasoning also responded during spatial attentional demand, and the size of these two effects (physical > social; hard > easy) were equally large. Third, individual differences in preferential engagement of physical > social and hard > easy fROIs in one task was predictable from individual differences in the other task. And fourth, responses from physical > social fROIs (but not hard > easy fROIs, and not a control region) predicted spatial working memory performance. Together, these findings suggest that neural resources for physical (vs social) reasoning are involved in functions beyond perceiving and reasoning about the physical world.

## 2. Materials and Methods

### 2.1 Open Science Practices

Some of our analyses were pre-registered and others were not; we mark these analyses as confirmatory and exploratory, respectively throughout the paper. Our pre-registration document can be found at https://osf.io/me8sc/. All data and code required to reproduce the results of this research can be found at https://osf.io/7mtk8.

### 2.2 Dataset

The dataset, including a total of 45 adult subjects, was collected by the last author (see Liu et al., 2024), and is openly available at https://openneuro.org/datasets/ds004934. We filtered this dataset to only include participants with 2 runs of data from a spatial working memory task (remembering more vs fewer locations in a grid), and 2 runs of data for a physical reasoning task (tracking and predicting the motions of two physical objects, versus two social agents). Our final sample was 28 adult participants (*M*_age_ = 26.5 years, 17 female, 50% White, 26 right-handed). These participants performed up to 2 other tasks, which are not the focus on the current work.

Neuroimaging data were collected from a 3-Tesla Siemens Magnetom Prisma scanner located at the Athinoula A. Martinos Imaging Center of the McGovern Institute, using the standard 32-channel head coil. Stimuli were projected to a screen behind the scanner, and were viewed by participants through a mirror mounted to the head coil, at a subtended visual angle of approximately 14 x 19 degrees. Each participant first underwent an anatomical scout scan (auto-align, acquired in 128 sagittal slices with 1.6 mm isotropic voxels; TA = 0.14; TR = 3.15 ms; FOV = 260 mm), followed by a high-resolution MPRAGE anatomical scan (T1-weighted structural images acquired in 176 interleaved sagittal slices with 1.0 mm isotropic voxels, TA = 5:53, TR = 2530.0 ms, FOV = 256 mm, GRAPPA parallel imaging, acceleration factor of 2). See Liu et al. (2024) for details. Afterwards, each participant completed two runs of the social vs physical prediction task, two runs of the spatial working memory task, which we describe below.

### 2.3 Physical prediction task

In the physical prediction task (Fischer et al., 2016), available at https://osf.io/sa7jy/, participants saw an overhead view of an arena containing two moving dots, and tracked the trajectories of the dots over the course of a 10 second video followed by a 1.5 second response window. In the physical condition, the dots interacted like billiard balls, bouncing off the walls of the arena and each other. In the social condition, the dots behaved like social agents, chasing and imitating each other. At the 7 second mark of the video, one dot disappeared for 3s. Participants were asked to continue to track its motion during these 3 seconds. At the end of this period, the dot reappeared, and subjects had 1.5s to judge whether it reappeared in a feasible or infeasible location via button press. Participants did not receive feedback on their performance. A 1.5s grey screen was displayed between each trial. This task was designed by Fischer et al. (2016) to be matched in difficulty across the social and physical conditions (and average performance was indeed equal across the two; see **SM Figure S1**). Each run of the task included 19 26-second blocks, each consisting of two 13-s trials; 8 social and 8 physical blocks; and 3 rest blocks, where participants saw a black screen for the full block duration. Each run lasted 8.2 minutes.

### 2.4 Spatial working memory task

In the spatial working memory task (Fedorenko et al., 2013), available at https://evlab.mit.edu/funcloc/, participants saw a white 8-by-8 grid. On each trial, participants saw a sequence of 4 events where cells in this grid changed from white to blue, and participants were asked to track the locations of these cells. In the hard condition, two cells in the grid changed at a time (adding up to 8 locations total), while in the easy condition, one cell changed at a time (adding up to 4 locations total). At the end of each trial, participants saw two alternative grids, one of which matched the sequence they saw, and indicated the correct grid via button press. Each trial was preceded by a fixation cross for 500 ms, followed by four grid patterns each displayed for 1000 ms, the two options of concatenated grids for 3750 seconds, and feedback for 250 ms. Each run included 20 16 second blocks (6 easy, 6 hard, and 4 rest blocks). Each run lasted 7.5 minutes.

### 2.5 Preprocessing and modeling

Data was preprocessed using fMRIprep (Esteban et al., 2019). This involved 3D motion correction, slice scan time correction, high-pass filtering of the BOLD signal (using a discrete cosine filter with 128s cut-off), motion correction using mcflirt (FSL v5.0.9), and volume-based spatial normalization to a standardized coordinate system (MNI152NLin6Asym; using Lanczos interpolation). The following confounds were estimated from the time-series of data each run: framewise displacement, derivative of variance over frame-to-frame motion, and global signals within each of the CSF, WM, and whole-brain masks. Volumes were flagged as motion outliers if framewise displacement exceeded 0.5 mm or if the derivative of variance exceeded 1.5. A longer description of the preprocessing parameters are available in the **SM Section 2**. We inspected the report generated by fMRIprep on the anatomical and functional data for each subject for quality assurance. For first level modeling, we used the Nipype library (Gorgolewski et al., 2011) to fit the run-level GLM over the output of preprocessed BOLD images with a smoothing kernel of 6mm, including all of the confounds estimated during pre-processing as nuisance regressors. We defined event regressors using a boxcar for the start and end of each task block (hard, easy, social, physical, rest). These event regressors were convolved with a double-gamma hemodynamic response function and a high-pass filter of 1/210 Hz. For second level modeling, we used the FSL FEAT sub-tool (within the Nipype package), FLAME (FMRIB’s Local Analysis of Mixed Effects) to create a single second-level model, per subject and per contrast.

### 2.6 Parcels and functional regions of interest

We defined 8 parcels (search spaces) to constrain the locations of fROIs. We combined frontal and parietal parcels for the multiple demand network (Fedorenko et al., 2013; Pramod et al., 2022) (https://evlab.mit.edu/funcloc), and parcels for regions preferentially engaged by physical reasoning (Fedorenko et al., 2013; Pramod et al., 2022). The resulting 8 parcels are shown in **Figure 1C**. These 8 parcels were then used to identify two sets of fROIs per participant: one set engaged by the spatial working memory task (multiple demand, MD, or hard > easy fROIs), and one set engaged by the physical processing task (intuitive physics,IP, or physical > social fROIs). For each parcel, participant, task, and run, we selected an fROI by taking the top 10% most active voxels from the *z* map contrasting hard > easy for the spatial working memory task, and physical > social for the physical prediction task. This procedure ensures that each fROI is the same size across participants, and can be found in every single participant; see **SM Section 8** for results from other top N% fROI definitions, and **SM Section 4.3** from an alternative approach that defines fROIs as suprathreshold voxels instead. The result of this procedure was 32 fROIs for each subject for each set of regions (IP and MD; 8 parcels, 2 tasks, 2 runs each).

**Figure 1.**
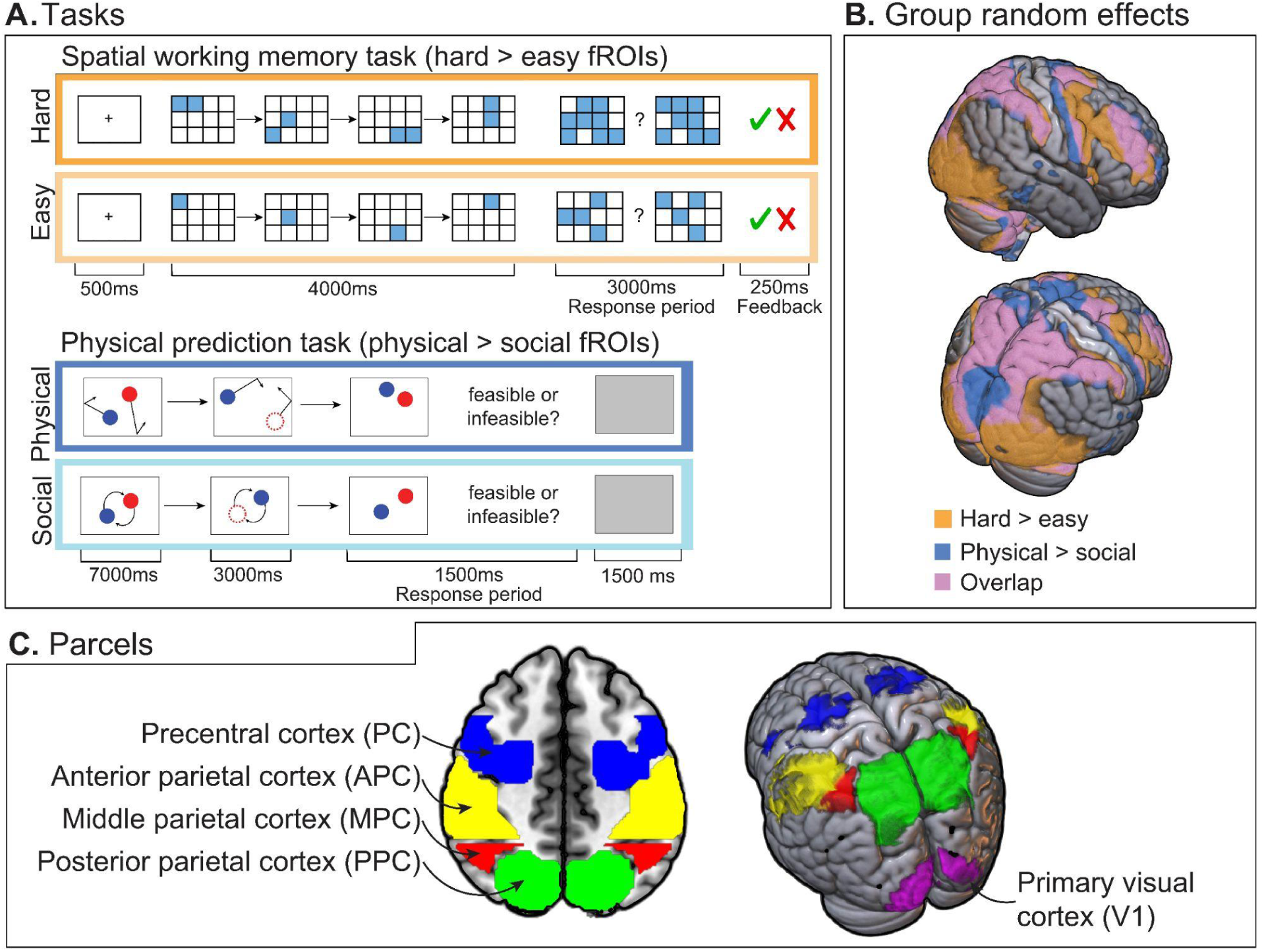
Overview of tasks and brain regions. (A) Conditions and trial structure of the spatial working memory and physical prediction tasks. (B) Results of a group random effects analysis (N = 28) mapping preferential responses to hard over easy blocks and physical over social blocks (nonparametric one-sample t-test with 5000 permutations, *α* = 0.05, one-tailed, using threshold-free cluster enhancement (TCFE) and variance smoothing of 6 mm. (C) Cortical parcels in frontal and parietal cortex, as well a control region, primary visual cortex (V1).

**Figure 2.**
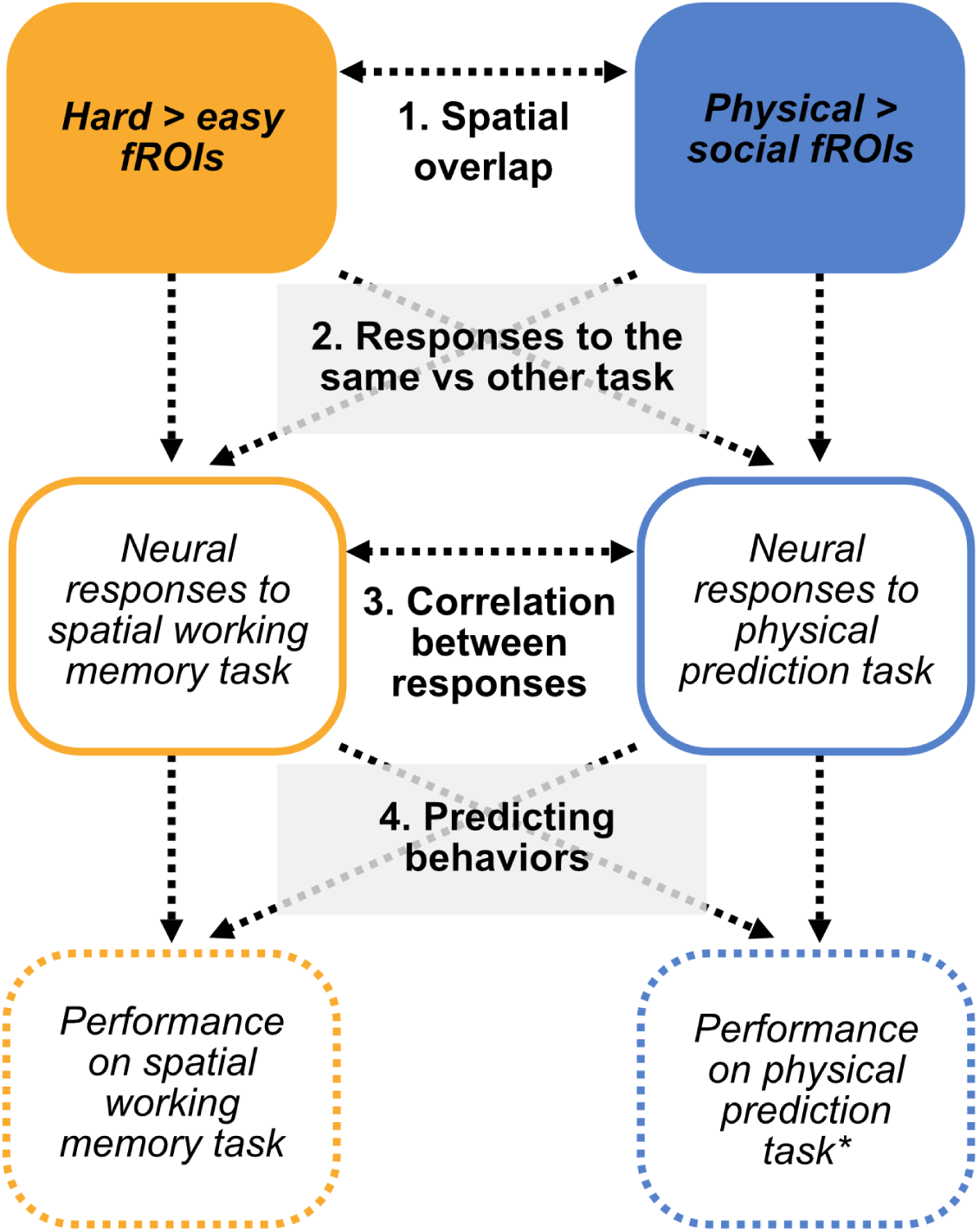
Overview of analyses. After identifying fROIs maximally engaged by the spatial working memory task and physical prediction task (top row), we (1) measured the spatial overlap between these fROIs, (2) measured their responses in held out data from the same task and opposite task, (3) related responses from fROIs in the same vs opposite set to each other, and (4) used neural responses to predict behavioral performance on each task (bottom row). *Because behavioral performance on the physical prediction task was unreliable across runs, we did not study the relationship between neural responses and behavioral performance on this task.

This allowed us to study responses of each fROI set held out data from the same task, as well as the opposite task. After pre-registering these parcels, we decided to add a control fROI, primary visual cortex (V1), which we identified by taking the top 10% most active voxels responsive to stimulus > rest in the spatial working memory task within an anatomically defined bilateral parcel. All parcels are available at OSF (https://osf.io/7mtk8).

### 2.7 Overview of fROI analyses

Given these two sets of fROIs, we measured (1) the degree of spatial overlap between hard > easy and physical > social fROIs in each parcel, (2) univariate responses in hard > easy fROIs to the physical vs social task (and vice versa), (3) correlations between univariate responses within and across the two sets of fROIs, and (4) relationships between neural responses in these fROIs and behavioral performance. In the analyses described below, all mixed effects models were implemented using the lme4 (Bates et al., 2015) and lmerTest (Kuznetsova et al., 2017) packages in R. Model formulas, including specification of fixed and random effects, are listed in each section below. Quality assurance of statistical models was conducted using the `check_models()` function from the performance package (Lüdecke et al., 2021).

#### 2.7.1 Spatial overlap (confirmatory)

How much do voxels maximally engaged by physical (vs social) processing, and those maximally engaged by hard (vs easy) spatial working memory, overlap? To answer this question, we computed the Dice’s Coefficient (DC) between each person’s physical > social and hard > easy fROI within each parcel. DC is defined by twice the number of overlapping voxels present in both fROIs divided by the total number of voxels across fROIs. In a first pre-registered analysis, we computed Dice’s Coefficient for pairs of fROIs in each parcel computed using second-level maps, folding across the two runs per task. In an exploratory analysis, we compared overlap between each person’s fROIs estimated from separate runs of the same task (e.g. hard > easy vs hard > easy) and across different tasks (hard > easy and physical > social).

#### 2.7.2 Univariate responses of fROIs across tasks (confirmatory)

In the frontal and parietal cortex, is activity in IP fROIs modulated by task difficulty, and are MD fROIs more responsive during physical than social processing? To address this question, we pre-registered an analysis of the profile of responses for each fROI across both tasks by modeling an interaction between task (physical prediction vs spatial working memory) and preference (hard > easy; physical > social). To ensure that the same amount of data contributed to analyses of responses within vs across tasks, we always studied neural responses across run numbers (i.e. hard > easy across runs 1-2; hard > easy run 1 vs physical > social run 2; physical social run 1 vs hard > easy run 2). Our model formula was: `beta ∼ preference * task + (1|subject)`.^1^ We used `lsmeans()` to extract the two pairwise comparisons from this interaction.

We also conducted additional exploratory analyses. The first repeats the confirmatory analysis over responses from all fROIs in each set, to examine the overall responses from hard > easy fROIs and physical > social fROIs (model formula `beta ∼ preference * task + (1|fROI) + (1|subject)`). The second compared the magnitude of hard > easy (or physical > social) responses across the two types of fROIs within the same anatomical parcel (`beta ∼ condition * fROI_set + (1|subject)`). If these two sets of fROIs constitute (at least partially) separate systems, then we should find that the spatial working memory task evokes a stronger hard > easy response in hard > easy fROIs, and that the physical prediction task evokes a stronger physical > social response in the physical > social fROIs. Further exploratory analyses in this line are described in the results.

#### 2.7.3 Correlations within and across fROI sets (exploratory)

Do univariate responses in IP fROIs predict univariate responses in MD fROIs? To answer this question, we studied individual differences in these two contrasts (physical > social and hard > easy) across vs within the two sets of fROIs in a series of exploratory analyses. If the two sets of fROIs are dissociable in function, then activity between any pair of fROIs should be more correlated when both regions come from the same set of regions, than when they come from different sets. If the two sets of fROIs are related in function, then activity of a given fROI during one task should predict its response in the second task. We computed Spearman’s rho (*ρ*) for each fROI to all other fROIs, excluding pairs of fROIs within the same parcel to prevent spatial overlap from contributing to results. This allowed us to study (1) the relationship between neural responses in IP fROIs and other IP fROIs, (2) between MD and other MD fROIs, and (3) between fROI sets.

We compared correlation strength with vs across fROI sets using a linear model (`correlation ∼ fROI_pair,` where fROI_pair indicated whether the two regions contributing to that correlation came from the same or different set; IP-MD, IP-IP, MD-MD). By setting IP-MD as the reference level of this predictor, we could compare the correlation between fROI sets (IP-MD) to the correlation within fROI sets (IP-IP and MD-MD). We used a permutation test to assess whether the correlation between fROIs across sets (i.e. IP-MD; the intercept of the model reported above) was significantly greater than 0. Specifically, we simulated a distribution of *t* statistics for the same intercept under the null hypothesis by shuffling participant labels within each fROI set, and fitting the same model, 5000 times. Then we computed the *p*-value of our empirical test statistic by calculating the proportion of *t* statistics from the null distribution that were equal to or greater than the observed empirical *t* statistic.

We also compared the degree of correlation between hard > easy and physical > social fROIs (IP-MD) to the correlation between both of these fROIs to a control fROI, V1 (V1-IP, V1-MD), in a separate model.

#### 2.7.4 Relating neural signals to behavior (exploratory)

How do neural responses from IP and MD fROIs relate to behavioral performance? Before exploring this question, we first checked whether these potential individual difference measures were reliable by computing the split-half reliability (Spearman’s *ρ*) of behavioral performance (% accurate), and neural responses (hard > easy; physical > social). For these exploratory analyses, we omitted one participant who lacked behavioral data for their spatial working memory task (N = 27). We also computed the correlation between regions, to evaluate whether neural responses and DC values could be averaged into a single predictor per set to simplify our subsequent analyses. We then conducted stepwise regressions to evaluate the neural predictors of individual differences in behavior, and a mediation analysis that tests how spatial overlap between fROIs contributes to this relationship, described further in the results.

## 3. Results

### 3.1 Task difficulty and reliability of measures (descriptive statistics)

#### 3.1.1 Task difficulty

Our tasks were well-matched for difficulty. Of the four conditions from our two tasks (physical and social prediction, hard and easy spatial working memory), people achieved an average of ∼80% correct on 3 of the 4 conditions, except for the easy condition of the spatial working memory task, for which average performance was 95% (see **Table 1**)^2^. Performance in the easy condition of the spatial working memory task differed from the three others (*p* < .001, two-tailed), and none of the other conditions differed from each other (**SM Figure S1**).

**Table 1.** Metrics describing task difficulty, and reliability of behavioral and neural individual differences. Reliability of neural measures (Spearman’s *ρ*, Spearman-Brown corrected) was computed over data averaged across runs and fROIs. Bolded values indicate measures that were reliable by a threshold of *α* = .05, two-tailed

| Task | Spatial working memory |  | Physical prediction |  |
| --- | --- | --- | --- | --- |
| Condition | Hard | Easy | Physical | Social |
| Accuracy $M$ (SD) | 80.3% (9.78) | 94.7% (7.55) | 80.0% (6.76) | 81.1% (10.3) |
| Reliability of performance $\rho$ ($p$) | $\rho = 0.71$, $p = \mathbf{0.003}$ | $\rho = 0.30$, $p = 0.370$ | $\rho = 0.22$, $p = 0.550$ | $\rho = 0.61$, $p = \mathbf{0.021}$ |
| Reliability of univariate preference averaged across fROIs within each set $\rho$ ($p$) | Hard > easy responses in:<br>• Hard > easy fROIs: $\rho = 0.89$, $p < \mathbf{0.001}$<br>• Physical > social fROIs: $\rho = 0.70$, $p < \mathbf{0.001}$ | | Physical > social responses in:<br>• Physical > social fROIs: $\rho = 0.95$, $p < \mathbf{0.001}$<br>• Hard > easy fROIs: $\rho = 0.79$, $p = \mathbf{0.003}$ | |

#### 3.1.2 Reliability of neural and behavioral data

Individual differences in spatial overlap between hard > easy and physical > social fROIs were reliable (DC between fROIs within an anatomical region, averaged across regions: *ρ* = 0.56, *p* = 0.041, two-tailed, Spearman-Brown corrected). Spatial overlap in these two sets of fROIs were also correlated across anatomical regions (*M_ρ_*= 0.52, range = 0.23, 0.74; **SM Figure S2**), such that people with high overlap in some parcels also tended to have high overlap in others. Therefore, we used average DCs across anatomical parcels in individual difference analyses relating neural data to behavioral performance.

Individual differences in all neural responses were reliable across runs (hard > easy and physical > social for IP and MD fROIs shown in **Table 1**; V1 stimulus > rest split-half *ρ* = 0.94, *p* < 0.001, two-tailed, Spearman-Brown corrected). Neural responses within each fROI set were highly correlated to each other (for hard > easy fROIs: *M_ρ_* = 0.73, range = 0.62, 0.90; for physical > social fROIs: *M_ρ_* = 0.68, range = 0.44, 0.83). Therefore, we averaged neural responses per fROI set for relating neural data to behavioral performance.

By contrast, the reliability of the behavioral data was uneven. We found reliable performance on the hard condition of the spatial working memory task (*ρ* = 0.71, *p* = 0.003, two-tailed, Spearman-Brown corrected), though not the easy condition (*ρ* = 0.30, *p* = 0.370, two-tailed, Spearman-Brown corrected), likely due to a ceiling effect. In the physical vs social prediction task, behavioral performance in the social condition (*ρ* = 0.61, *p* = 0.021, two-tailed,

Spearman-Brown corrected) was reliable, but not the physical condition (*ρ* = 0.22, *p* = 0.550, two-tailed, Spearman-Brown corrected). Thus, for our research question concerning physical understanding and spatial working memory in the frontoparietal cortex, only the hard condition of the spatial working memory task was well suited for individual difference analyses.

### 3.2 IP and MD fROIs moderately overlap (confirmatory)

On average, the Dice’s Coefficient between physical > social and hard > easy second-level fROIs within each parcel was 0.24 (range = 0.22, 0.28) (**Figure 3a**, see **SM Figure S11** for how this scales with fROI size). Using the heuristics from Wilson et al. (2017), this would indicate a small degree of overlap. However, this assumes 100% overlap is possible in our data. To investigate this further, in an exploratory analysis we compared the overlap in fROIs derived from a single run of the two tasks with overlap in fROIs derived from two runs of the same task. We could then treat the overlap in within-task fROIs as a noise ceiling: how much overlap we should expect at maximum given our data. We found that the average DC between physical > social and hard > easy first-level fROIs was 0.17 (range = 0.15, 0.21) (**Figure 3b**). However, the maximum DC we expect given overlap between fROIs from the same task was on average 0.43 (range = 0.28, 0.55). By scaling the across-task DC values within each parcel by this noise ceiling, we found an average “adjusted” DC of 0.49 (range = 0.34, 0.74; **SM Table S1**).

**Figure 3.**
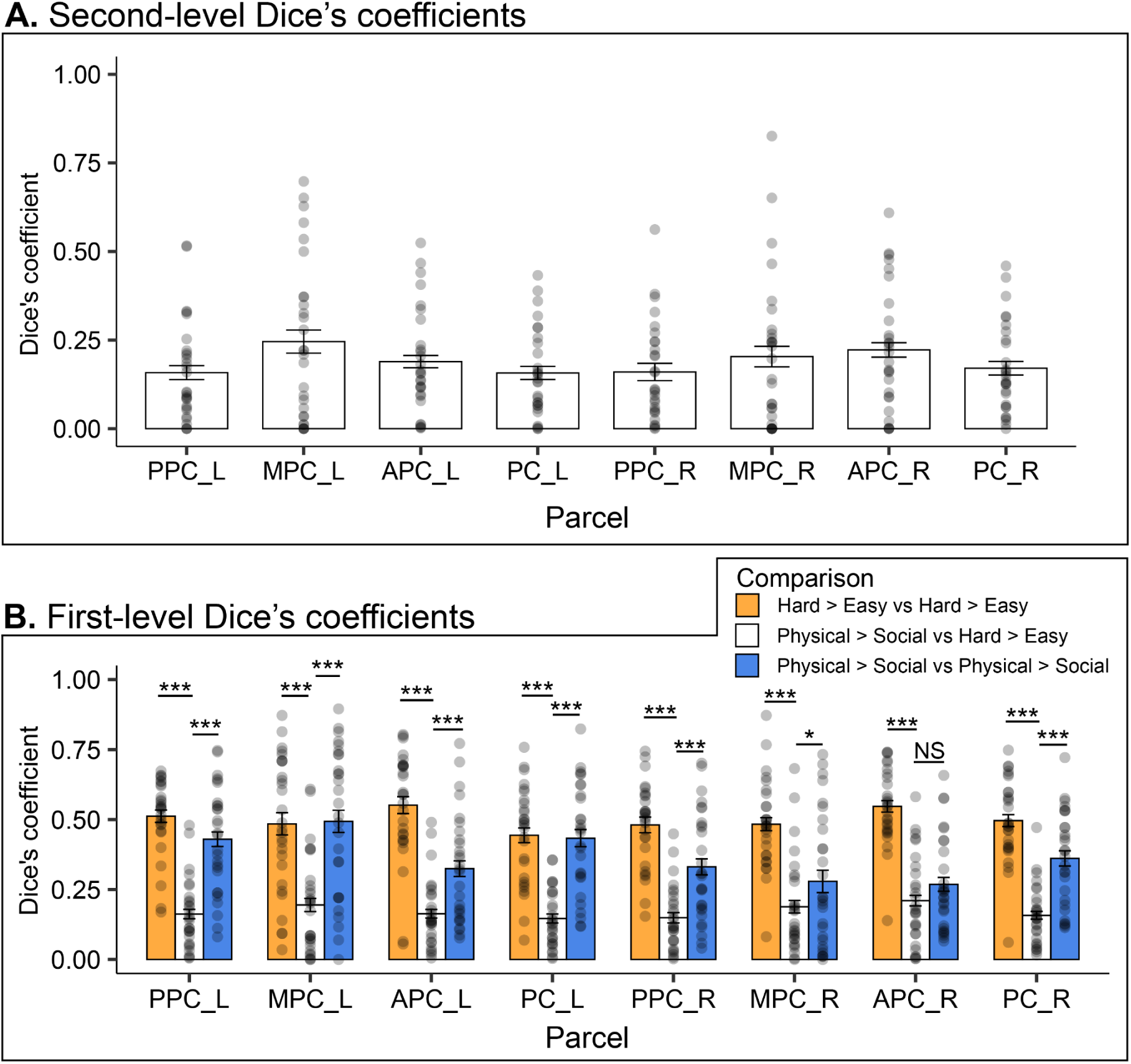
Overlap in fROIs within and across tasks. (A) Dice’s coefficient between physical > social fROIs and hard > easy fROIs for each anatomical parcel. (B) Dice’s coefficient between fROIs estimated in two partitions of data from the same task (blue and orange) and between the two tasks (white). NS indicates *p* >= .1; + < .1; * < .05; **< .01; *** < .001, two-tailed. Error bars indicate standard error of the mean.

In a separate exploratory analysis, we measured the spatial extent of voxels in each parcel that preferentially responded to the hard > easy, and physical > social contrast. First, we summed up the number of suprathreshold voxels (*p* < .001, uncorrected) for each person for each parcel for each contrast. We found that the number of suprathreshold hard > easy voxels was much greater (∼2-4x greater) than the number of physical > social voxels in the same parcel (**SM Figure S3**). This indicates that the spatial working memory task evokes a broader set of responses in the frontoparietal cortex than the physical prediction task in individual participants.

Is the physical > social contrast selecting a spatially restricted sub-population within a broader set of regions responsive to general task demands? In exploratory analyses addressing this question, we found evidence against this possibility. First, we found that the majority of physical > social suprathreshold voxels (*p* < .001, uncorrected) did not overlap with hard > easy voxels at the same threshold. Only 30.2% physical > social voxels (averaged across parcels; range 19.2-45.7%) responded significantly more to hard > easy. Likewise, only 13.9% hard > easy voxels (range 7.6%-22.1%) responded significantly to physical > social (see **SM Figure S4**). Looking at just the top 10% fROIs, 22.9% of voxels in IP fROIs (averaged across parcels: range 13.1-42.6%) were also responsive to spatial demand (*p* < .001, uncorrected), and 6.4% of voxels in MD fROIs (range 3.7-11.3%) were responsive to physical > social (*p* < .001, uncorrected). Thus, the hard > easy contrast evokes a broader set of responses in the frontoparietal cortex than the physical > social contrast, but the physical > social contrast still picks out partially distinct voxels that are not merely a subset of this broader hard > easy response. The asymmetry of these two responses will mirror results in following sections.

As a last exploratory analysis characterizing the spatial topography of responses from these two tasks, we found that hard > easy responses were right-lateralized, and physical > social responses left-lateralized. We computed a lateralization index using the formula LI = (LV - RV) / (LV+ RV) (Brumer et al., 2020) where LV and RV are the number of suprathreshold voxels (*p* < 0.001, uncorrected) per person across all 4 anatomical parcels per hemisphere. Physical > social responses were significantly above 0 (left-lateralized; *B* = 0.49, 95% CI = [0.35, 0.63], *p* < 0.001, two-tailed), hard > easy response were significantly below 0 (right-lateralized; *B* = −0.24, 95% CI = [−0.10, −0.38], *p* = 0.001, two-tailed) (difference in LIs, *B* = 0.73, 95% CI = [0.53, 0.93], *p* < 0.001, two-tailed). See **SM Section 3.3** for details. This mirrors prior work by Osiurak and colleagues (2024), who reported a region in left IPL that was responsive during physical reasoning but not Raven’s matrices, and in prior work on deficits in tool use and mechanical reasoning that often results in left-hemisphere lesions in parietal cortex (Goldenberg & Hagmann, 1998; but see Liu et al., 2025).

### 3.3 Activity in IP fROIs is modulated by spatial attentional demand (confirmatory)

The univariate responses from both sets of fROIs are shown in **Figure 4** and **Table 2**. All 8 hard > easy fROIs and 7 out of 8 physical > social fROIs responded reliably to the same contrasts in held out data.

**Figure 4.**
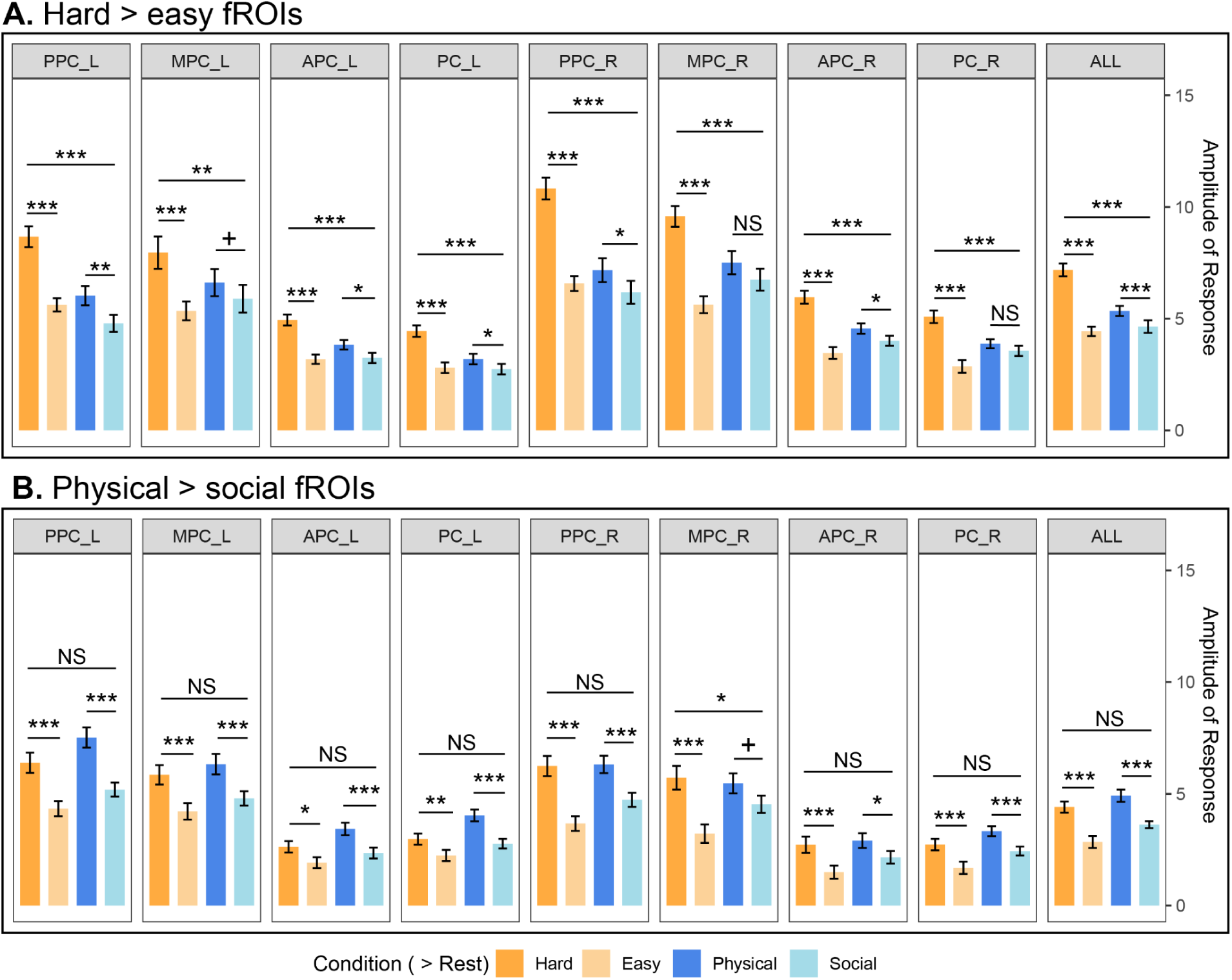
Responses in (A) hard > easy fROIs and (B) physical > social fROIs during hard vs easy spatial working memory (left; in orange) and physical vs social prediction (right; in blue). All 8 hard > easy fROIs showed a greater hard > easy than physical > social response, but no physical > social fROI showed a greater physical > social than hard > easy response. NS indicates *p* >= .1; + < .1; * < .05; **< .01; *** < .001, two-tailed. Error bars indicate standard errors. See also Table 2.

**Table 2.** Univariate responses to both tests in hard > easy (multiple demand, MD) and physical > social (intuitive physics, IP) fROIs. Significant *p*-values (threshold of *α* = .05, two-tailed) are bolded. * indicates significant *p-*values from individual regions that pass a more conservative corrected threshold of *α* = .00625 (8 fROIs per ROI set, MD and IP).

| fROI | $df$ | Physical prediction > social prediction | | Hard > easy spatial working memory | | Interaction | |
| --- | --- | --- | --- | --- | --- | --- | --- |
| | | Std. $\beta$ | $p$-value | Std. $\beta$ | $p$-value | Std. $\beta$ | $p$-value |
| MD PPC_L | 193 | 0.29 | **0.005\*** | 0.72 | **<0.001\*** | 0.43 | **0.003\*** |
| MD MPC_L | 193 | 0.13 | 0.093 | 0.45 | **<0.001\*** | 0.33 | **0.002\*** |
| MD APC_L | 193 | 0.23 | **0.016** | 0.69 | **<0.001\*** | 0.46 | **0.001\*** |
| MD PC_L | 193 | 0.21 | **0.035** | 0.78 | **<0.001\*** | 0.56 | **<0.001\*** |
| MD PPC_R | 193 | 0.21 | **0.049** | 0.88 | **<0.001\*** | 0.67 | **<0.001\*** |
| MD MPC_R | 193 | 0.16 | 0.132 | 0.84 | **<0.001\*** | 0.68 | **<0.001\*** |
| MD APC_R | 193 | 0.22 | **0.047** | 1.00 | **<0.001\*** | 0.78 | **<0.001\*** |
| MD PC_R | 193 | 0.15 | 0.209 | 1.07 | **<0.001\*** | 0.91 | **<0.001\*** |
| MD all fROIs | 1754 | 0.17 | **<0.001** | 0.66 | **<0.001** | 0.49 | **<0.001** |
| IP PPC_L | 193 | 0.56 | **<0.001\*** | 0.49 | **<0.001\*** | 0.07 | 0.649 |
| IP MPC_L | 193 | 0.38 | **<0.001\*** | 0.41 | **<0.001\*** | -0.03 | 0.840 |
| IP APC_L | 193 | 0.45 | **<0.001\*** | 0.30 | **0.015** | 0.15 | 0.374 |
| IP PC_L | 193 | 0.58 | **<0.001\*** | 0.33 | **0.002\*** | 0.24 | 0.105 |
| IP PPC_R | 193 | 0.40 | **<0.001\*** | 0.65 | **<0.001\*** | -0.25 | 0.116 |
| IP MPC_R | 193 | 0.24 | 0.076 | 0.63 | **<0.001\*** | -0.40 | **0.036** |
| IP APC_R | 193 | 0.30 | **0.021** | 0.49 | **<0.001\*** | -0.19 | 0.298 |
| IP PC_R | 193 | 0.45 | **<0.001\*** | 0.53 | **<0.001\*** | -0.08 | 0.655 |
| IP all fROIs | 1754 | 0.37 | **<0.001** | 0.44 | **<0.001\*** | -0.07 | 0.250 |

How do these fROIs respond during the opposite task? We found that whereas MD fROIs were more tuned to task demands than physical content, IP fROIs responded just as strongly to hard > easy spatial working memory as they did to physical > social content. While 5 out of 8 MD fROIs responded more during physical than social prediction, only 1 fROI met a more stringent significance threshold correcting for the 8 regions in this set. All 8 MD fROIs showed a significant task-by-condition interaction, such that they showed a bigger hard > easy than physical > social response. By contrast, all 8 IP fROIs (7 meeting a more stringent significance threshold) responded more to the hard than the easy condition of the spatial working memory task. Only one IP fROI, right MPC, showed a significant task-by-condition interaction, but in the opposite direction than anticipated (i.e. greater hard > easy than physical > social responses).

Folding across all regions within each fROI set, we found a significant task-by-condition interaction in MD fROIs, with greater responses to hard > easy than physical > social blocks, but no such interaction in IP fROIs, replicating Kean et al. (2025). See **Table 2**. These findings were stable across a range of fROI sizes (top 1%-50%; see **SM Figure S12**), and in top 10% fROIs removing overlapping voxels between the two contrasts (**SM Section 4.1**). We again found this overall pattern when we chose fROIs not by taking the top N%, but by including all suprathreshold voxels (*p* < 0.001, uncorrected) per region per person (**SM Section 4.3**).

In a further exploratory analysis, we found a significant neural response-by-fROI set interaction, such that hard > easy responses were larger in the MD regions than in IP regions (*β* = 0.30, 95% CI = [0.17, 0.43], *p* < 0.001, two-tailed), and vice-versa where physical > social responses were larger in IP regions than in MD regions (*β* = 0.15, 95% CI = [0.02, 0.28], *p* = 0.020, two-tailed). This finding replicates Kean et al. (2025). However, this result only appeared when folding together all 8 regions within each set; looking at fROIs within each anatomical region separately, we observed greater hard > easy responses in MD than IP fROIs in 5 anatomical regions (2 meeting a more stringent threshold), and a greater physical > social response in IP than MD fROIs in 1 anatomical region (0 meeting a more stringent threshold); see **SM Table S2**.

Thus, voxels in the frontal and parietal cortices most engaged by a difficult (vs easy) spatial working memory task were not strongly tuned to physical (over social) processing, but voxels in the same cortical territory most engaged by physical (vs social) processing were also responsive to general attentional demands. Together, these results provide evidence both for dissociability of physical inference and attentional demand (since there are portions of frontoparietal cortex that is only responsive for attentional demand), and for the connection between these two mental functions (since there are also portions of frontoparietal cortex responsive to both contrasts).

### 3.4 Responses in MD fROIs predict responses in IP fROIs (exploratory)

Do individual differences in MD fROIs predict individual differences in IP fROIs? First, we measured the correlation of neural responses between vs within fROI sets during just the physical prediction task (physical > social), and during just the spatial working memory task (hard > easy). We found that for both tasks, the correlation of responses within an fROI set was higher than across set (during the physical prediction task: *B*_IP-IP_ = 0.16, 95% CI = [0.11, 0.21], *p* < 0.001, two-tailed; *B*_MD-MD_ = 0.15, 95% CI = [0.10, 0.21], *p* < 0.001, two-tailed; during the spatial working memory task: *B*_IP-IP_ = 0.13, 95% CI = [0.07, 0.18], *p* < 0.001, two-tailed; *B*_MD-MD_ = 0.17, 95% CI = [0.12, 0.23], *p* < 0.001, two-tailed). Yet, at the same time, for both tasks, correlations between responses across fROI sets was still significantly higher than 0 (during the physical prediction task: *B*_IP-MD_ = 0.53, 95% CI = [0.50, 0.56], *p* < 0.001, one-tailed; during the spatial working memory task; *B*_IP-MD_ = 0.55, 95% CI = [0.53, 0.59], *p* < 0.001, one-tailed; *p*-values from permutation test against the null distribution achieved by shuffling participant labels). Responses between MD and IP fROIs were also more correlated to each other, than when compared to V1 (*B* = −1.09, 95% CI = [−1.60, −0.59], *p* < 0.001, two-tailed; **SM Table S5**), suggesting that this effect is specific to regions in frontoparietal cortex. See **SM Figure S9** for pairwise correlations between individual fROIs.

Next, we tested whether people with large hard > easy responses in MD fROIs also tended to have large physical > social responses in IP fROIs. We found that although hard > easy responses from MD fROIs and physical > social responses from IP fROIs were more correlated to themselves than to each other (*B*_IP-IP_ = 0.42, 95% CI = [0.37, 0.47], *p* < 0.001, two-tailed; *B*_MD-MD_ = 0.46, 95% CI = [0.41, 0.51], *p* < 0.001, two-tailed), these two distinct responses from these two sets of fROIs were still significantly correlated across tasks (*B*_IP-MD_ = 0.27, 95% CI = [0.24, 0.30], *p* < 0.001, one-tailed, permutation test). See **Figure 5C**. Again, the correlation between hard > easy and physical > social fROIs across tasks was greater than the correlation between these fROI sets and visual responses in V1 (*B* = 0.17, 95% CI = 0.09, 0.25], *p* < 0.001, two-tailed; **SM Table S5**). Thus, even across two distinct tasks, neural activity in IP regions predicts activity in MD regions (**Table 3**). We found similar results when we varied the size of the fROIs (**SM Figure S13**). Overall, these findings suggest that functions in these two sets of fROIs are related to each other: Participants with stronger hard > easy responses in MD fROIs tended to have stronger physical > social responses in IP fROIs. At the same time, these two functions are not redundant: the relationship between responses was larger within than across fROI sets.

**Figure 5.**
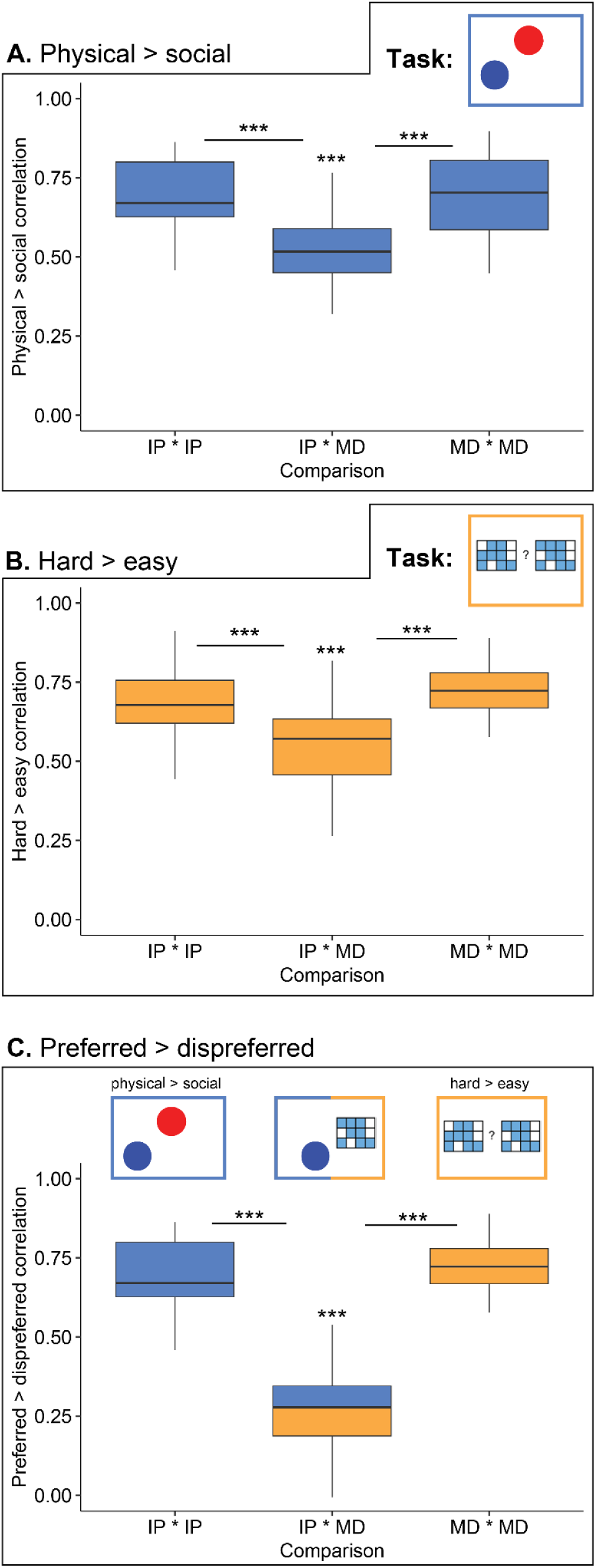
Relationship between hard > easy and physical > social responses, within and across fROI sets. Each panel shows the correlation (Spearman’s *ρ*) of preferential responses within IP fROIs, within MD fROIs and between these two fROI sets (IP-MD). fROIs within the same parcel were excluded from this analysis. NS indicates p >= .1; + < .1; * < .05; **< .01; *** < .001, two-tailed.

**Table 3.**
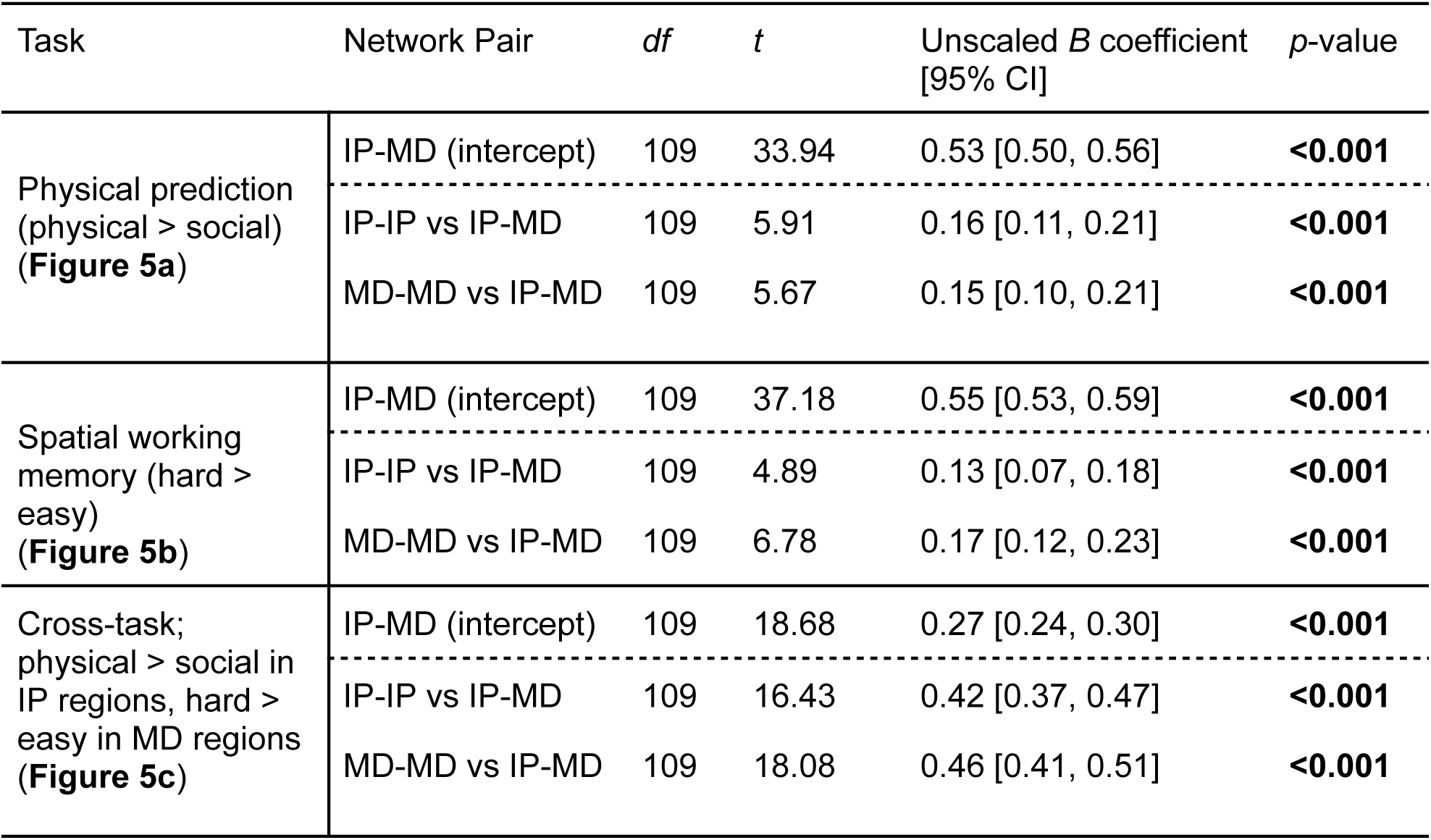
Correlation in univariate effects between vs within hard > easy (multiple demand, MD) and physical > social (intuitive physics, IP). Bolded values indicate significance (*p* < 0.05, two-tailed for comparing within to between-network values; one-tailed for comparing the between-network values to 0.

| Task | Network Pair | *df* | *t* | Unscaled *B* coefficient<br>[95% CI] | *p*-value |
| --- | --- | --- | --- | --- | --- |
| Physical prediction<br>(physical > social)<br>(**Figure 5a**) | IP-MD (intercept) | 109 | 33.94 | 0.53 [0.50, 0.56] | **<0.001** |
|  | IP-IP vs IP-MD | 109 | 5.91 | 0.16 [0.11, 0.21] | **<0.001** |
|  | MD-MD vs IP-MD | 109 | 5.67 | 0.15 [0.10, 0.21] | **<0.001** |
| Spatial working<br>memory (hard ><br>easy)<br>(**Figure 5b**) | IP-MD (intercept) | 109 | 37.18 | 0.55 [0.53, 0.59] | **<0.001** |
|  | IP-IP vs IP-MD | 109 | 4.89 | 0.13 [0.07, 0.18] | **<0.001** |
|  | MD-MD vs IP-MD | 109 | 6.78 | 0.17 [0.12, 0.23] | **<0.001** |
| Cross-task;<br>physical > social in<br>IP regions, hard ><br>easy in MD regions<br>(**Figure 5c**) | IP-MD (intercept) | 109 | 18.68 | 0.27 [0.24, 0.30] | **<0.001** |
|  | IP-IP vs IP-MD | 109 | 16.43 | 0.42 [0.37, 0.47] | **<0.001** |
|  | MD-MD vs IP-MD | 109 | 18.08 | 0.46 [0.41, 0.51] | **<0.001** |

### 3.5 Spatial overlap between MD and IP fROIs predicts larger univariate effects, which in turn predict better spatial working memory (exploratory)

How do individual differences in spatial overlap between fROIs and responses in those fROIs relate to spatial working memory behavior? **Figure 6** shows zero-order correlations between all pairs of univariate contrasts (averaged across runs and across fROIs within a set), spatial overlap between fROIs (averaged across regions), and behavioral measures (averaged across runs).

**Figure 6.**
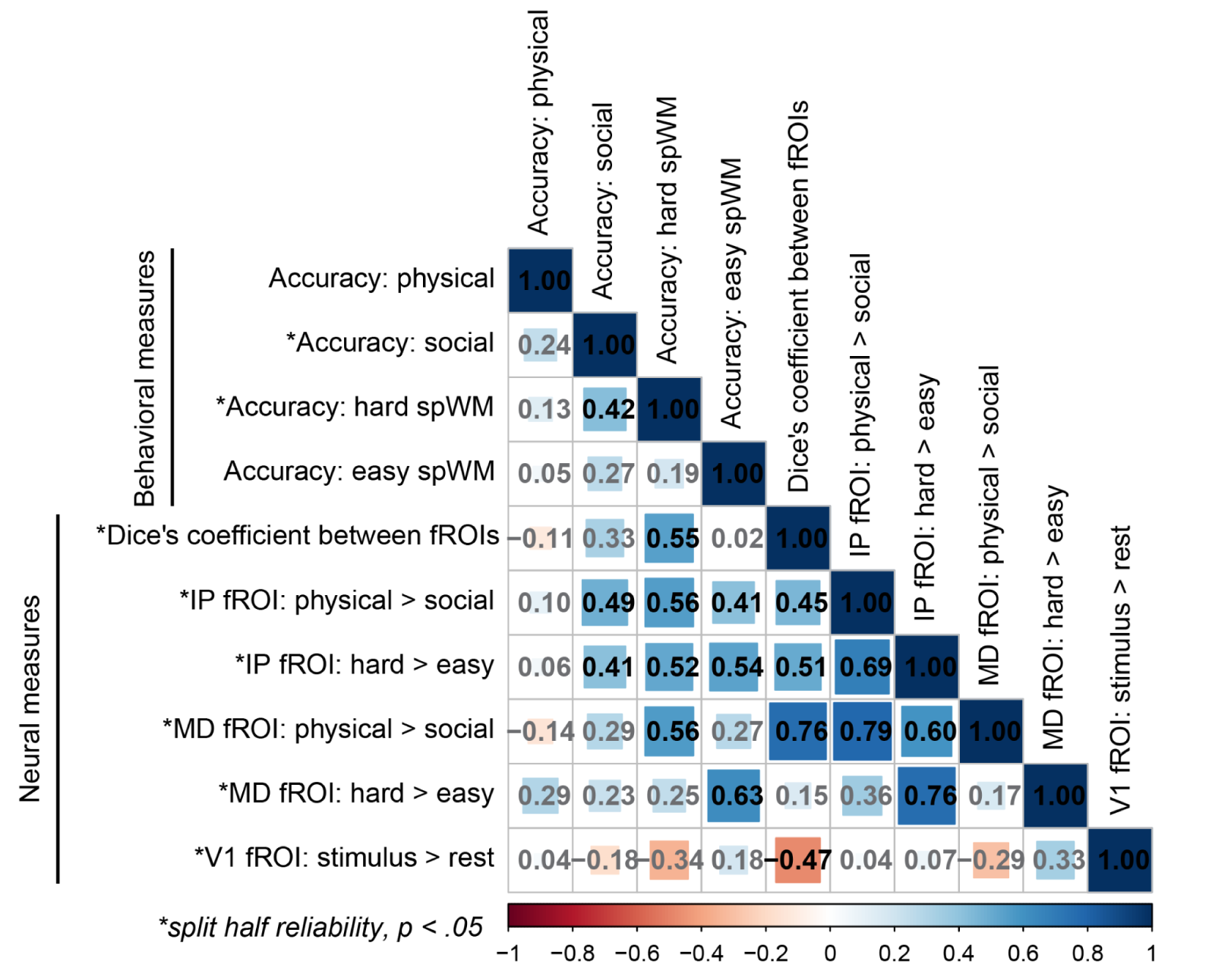
Spearman’s *ρ* between neural and behavioral measures across individuals. Accuracy was averaged across runs. Neural responses were averaged across runs and fROIs.* indicates reliable measures (*p* < 0.05, two-tailed). Insignificant correlations are shown in grey (*p* > 0.05, two-tailed).

People who were more accurate at the spatial working memory task also tended to have: (1) stronger hard > easy responses from IP (*ρ* = 0.519, *p* = 0.005, two-tailed) but not MD (*ρ* = 0.249, *p* = 0.211, two-tailed) fROIs; (2) stronger physical > social responses from both MD (*ρ* = 0.564, *p* = 0.002, two-tailed) and IP fROIs (*ρ* = 0.558, *p* = 0.002, two-tailed); and (3) greater overlap between MD and IP fROIs (*ρ* = 0.547, *p* = 0.003, two-tailed). By contrast, behavioral performance was not correlated with V1 responses. No measures were related to performance on the physical prediction task, though this is difficult to interpret, given the low reliability of the behavioral data from that task.

The main surprising positive finding from this analysis is that responses from IP fROIs predict spatial working memory performance. The main surprising negative finding is that responses from MD fROIs during the spatial working memory task did not predict performance on that very same task (*ρ* = 0.249, *p* = 0.211, two-tailed; **Figure 7**), despite the fact that data from hard > easy fROIs were highly reliable, and a prior paper reported a robust relationship between responses in these regions and performance on this same task (Assem et al. 2020), albeit with a much larger sample size.

**Figure 7.**
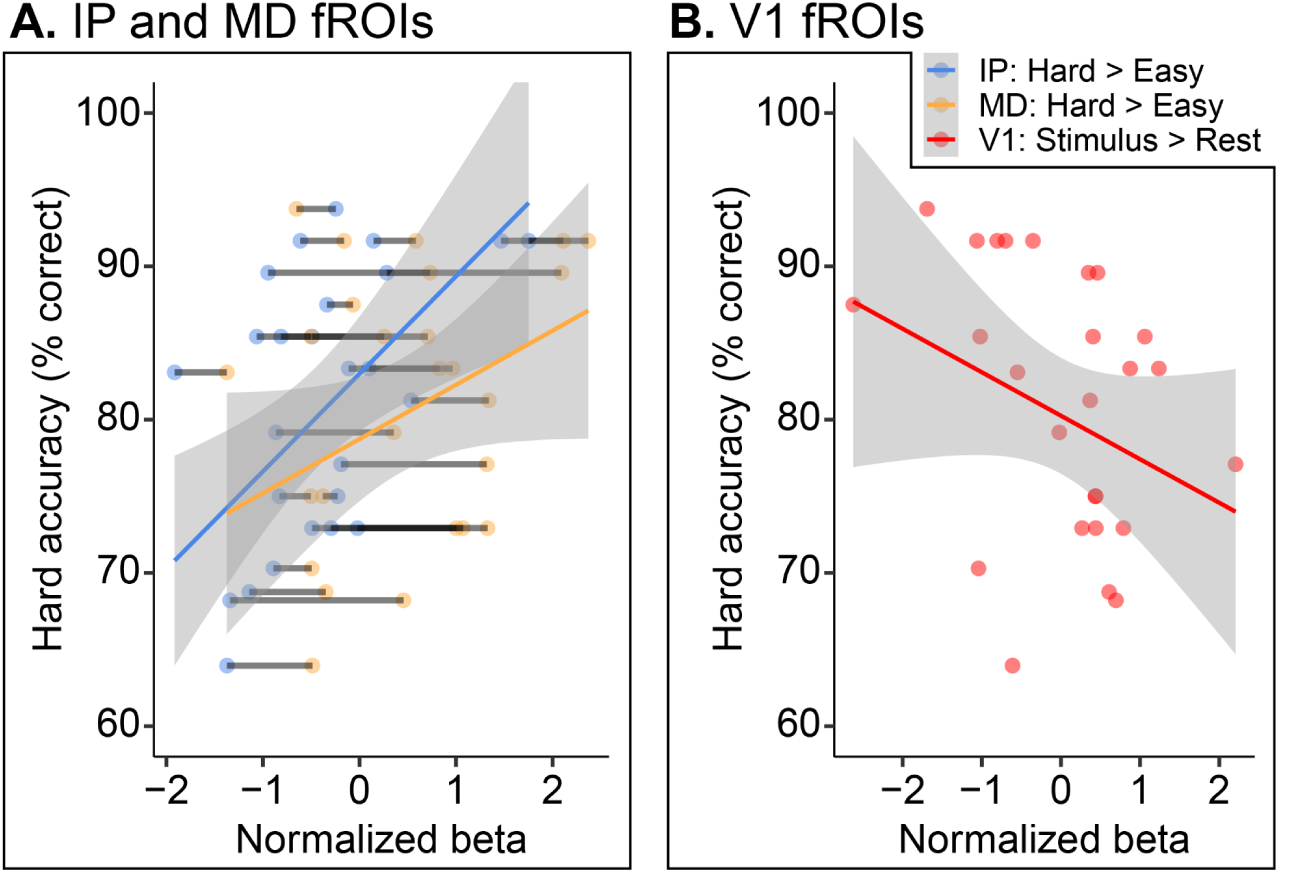
Relationships between individual differences in neural responses during the spatial working memory task, and performance during that task. (A) Hard > easy responses from MD and IP fROIs plotted against behavioral performance. (B) Stimulus > rest responses from V1 plotted against the same behavioral data. Behavioral performance was averaged across runs per subject. Neural responses were averaged across runs and fROIs, then normalized (e.g. scaled in terms of standard deviation away from the grand mean). Grey lines in (A) connect data from the same participants. Solid line and ribbon indicate the X and 95% confidence interval.

Our main question was which of these neural measures uniquely predicted spatial working memory performance. We conducted two sets of step-wise regressions to explore this question. First, we modeled performance as a function of fixed effects added to a linear regression in the following order: hard > easy responses from MD fROIs; hard > easy responses from IP fROIs; physical > social responses from MD fROIs; physical > social responses from IP fROIs; stimulus > rest response from V1. We chose this sequence by prioritizing neural signals during the same task, then from the opposite task, and then finally from the control fROI. At each step, we used a likelihood ratio test comparing each pair of models, as well as model AIC and BICs, to evaluate which set of predictors provided the best fit to the data without overfitting. Second, we trimmed this set of predictors to only those with significant zero-order correlations to behavioral performance (hard > easy responses from IP fROIs; physical > social responses from MD fROIs; physical > social responses from IP fROIs), and repeated the same steps. The full results of these analyses are reported in **SM Section 6**.

Both stepwise regressions showed that neural responses from physical > social fROIs explained variance in spatial working memory behavior (**Table 4**). When considering all potential neural predictors, the best model from our step-wise forward regressions included hard > easy responses from both sets of fROIs. In that model, hard > easy responses from IP fROIs (*β* = 0.77, 95% CI = [0.17, 1.37], *p* = 0.014, two-tailed) but not MD fROIs (*β* = −0.28, 95% CI = [−0.88, 0.32], *p* = 0.350, two-tailed) predicted behavior. However, hard > easy responses from MD fROIs did not uniquely explain variance in this model (likelihood ratio test, comparing model including both predictors with model including just hard > easy responses from IP fROIs: *F*(25) = 0.91, *p* = 0.350, see **SM Table S8**). When considering only neural predictors with positive zero-order correlations to behavior, the best model was one including only the hard > easy response from IP fROIs (*β* = 0.55, 95% CI = [0.20, 0.89], *p* = 0.003, two-tailed). These results were robust to fROI size (see **SM Figure S14**). They suggest that beyond merely responding during the spatial working memory task, hard > easy responses in physical > social fROIs predict individual differences in performance during that task.

**Table 4.** Regression results relating spatial working memory performance to neural activity using all possible predictors after stepwise model comparisons (see **SM Table S6**) and significant zero-order predictors after stepwise model comparisons (see **SM Table S7**). Bolded values indicate significance (*p* < 0.05).

| Model | fROI set | Response | df | t-value | Std. $\beta$ (95% CI) | p-value |
| --- | --- | --- | --- | --- | --- | --- |
| Best model, using all potential predictors | MD fROIs | Hard > easy | 24 | -0.953 | -0.28 [-0.88, 0.32] | 0.350 |
|  | IP fROIs | Hard > easy | 24 | 2.651 | 0.77 [0.17, 1.37] | **0.014** |
| Best model using only significant predictors | IP fROIs | Hard > easy | 25 | 3.253 | 0.55 [0.20, 0.89] | **0.003** |

Lastly, we explored a potential mechanism connecting spatial overlap between IP and MD fROIs, stronger hard > easy responses in IP fROIs, and enhanced spatial working memory performance: that people with more enmeshed responses (greater overlap between hard > easy and physical > social fROIs) had stronger peak responses in IP fROIs, which in turn predict stronger performance. (We did not test this with respect to responses in MD fROIs, which themselves did not predict performance.) We conducted an exploratory mediation analysis, implemented in the mediation package in R (Tingley et al., 2014). The relationship between better behavioral performance and stronger hard > easy responses in IP fROIs, was partially mediated by the overlap between MD and IP fROIs (*β* = 0.22, 95% CI = [0.01, 0.46], *p* = 0.046, two-tailed, average causal mediation effect). See **SM Section 7** for details.

## 4. Discussion

Our capacity to reason about the physical world involves representing and tracking objects, and making inferences and predictions about those objects (e.g. How heavy is this thing? What is it made of? Will it fall and in which direction?). Converging evidence from cognitive neuroscience shows that regions of the frontal and parietal cortices are preferentially involved in tasks designed to evoke physical reasoning (Fischer et al., 2016; Martin & Weisberg, 2003; Osiurak et al., 2025; Pramod et al., 2022), and contain representations that are distinctive to that domain ((e.g. mass, support, contact, containment; physical prediction error; Liu et al., 2024; Pramod, Mieczkowski, et al., 2025; Schwettmann et al., 2019). At the same time, work has shown that cortical regions in the same neighborhood are involved in domain-general attentional demand (Duncan, 2010). Prior work directly relating these two capacities in the same individual participants showed that partially dissociable regions are involved in physical reasoning and working memory (Kean et al., 2025), and that these behaviors are separable in individual differences (Mitko et al., 2024; Mitko & Fischer, 2020).

In this paper, we build on this prior research and compare neural responses in two difficulty-matched tasks: a task that contrasts tracking and making predictions about inanimate objects vs animate agents (Fischer et al., 2016), and a spatial working memory task that contrasts more vs less demanding engagement of spatial attention (Fedorenko et al., 2013).

Like Kean et al. (2025), who used the same spatial working memory task but a different unstable towers intuitive physics tasks, we found some degree of overlap in voxels maximally engaged by high (vs low) spatial working memory demand and by physical processing. Also like Kean et al. (2025), we found greater neural responses to hard than easy tasks in most physics fROIs and equal responses to physical and a control task in most multiple demand fROIs. This suggests that the asymmetry of the responses to spatial working memory demand in intuitive physics vs MD fROIs is robust to these two different physical reasoning tasks, which involve two different ways of defining physical reasoning (physical vs social prediction; physical prediction vs perceptual judgment). Unlike Osiurak et al. (2025), we did not find a left inferior parietal region that responded preferentially to a physical task but not a control task; instead, we found another region in the same anatomical area that showed the opposite preference, responding equally to physical and social processing while responding preferentially to general task demands.

Going beyond prior work, we also found robust individual differences in the preferential responses across fROI sets, even across tasks; individuals with a stronger hard > easy response during the spatial working memory task also showed a stronger physical > social response during the physical prediction task. These responses predicted behavior during spatial working memory: individuals with stronger activation in IP fROIs during the hard vs easy blocks of the spatial working memory task also were more accurate. (We were unable to study individual differences in the physical prediction task, because of low split-half reliability of the behavioral data.) From these findings, we suggest that portions of the frontal and parietal cortex preferentially responsive during physical reasoning are preferentially engaged and support performance during other tasks, outside of the domain of intuitive physics.

### 4.1 How domain-general are cortical regions engaged for processing of physical objects?

Our results have implications for the functional specificity of the so-called intuitive physics network. The intuitive physics network should be considered a specialized network insofar as it is more involved during physical reasoning than other tasks, contains information relevant to solving problems in the proposed task domain, is intrinsically connected to itself compared to nearby regions, and is causally involved in physical reasoning. So far, there is evidence that these regions are more connected to themselves than MD regions at rest (Pramod, Hutchinson, et al., 2025), contain information relevant to solving physical problems (e.g. stability, mass, contact) (Pramod et al., 2022; Pramod, Mieczkowski, et al., 2025; Schwettmann et al., 2019), and in group analyses, respond more to physical reasoning tasks than merely demanding alternatives (Fischer et al., 2016; Kean et al., 2025; Osiurak et al., 2025). Here, however, cortical regions that preferentially responded when people tracked and predicted movements of physical objects (vs social agents) also responded more when they tracked and remembered many (vs few) locations in a spatial grid; and the stronger these responses were, the better people were at the spatial working memory task. This is one piece of evidence against the strongest form of domain-specificity for the intuitive physics network, that this network is specialized for that specific mental function, but should be interpreted together with many potential sources of evidence.

Neural resources for physical reasoning appear to be involved in functions beyond this domain. What is the scope of these functions? First, it is possible that regions preferentially engaged during physical reasoning are very functionally broad. Although they do respond more to physical than social stimuli, and respond more during component processes of physical reasoning (e.g. prediction), it is possible that they respond to task difficulty more broadly and would support performance on any demanding task. Second, it is possible that these regions are tuned to task demands in the visuospatial domain in particular. In the current work, our working memory task requires representing and remembering locations, and reporting on that memory on every trial; the physical task also requires representing, predicting, and maintaining the position of physical objects in working memory while the object is under occlusion. Fedorenko et al. (2013) has already established that the very same voxels responded more to difficult than easy versions of many different tasks, including both verbal and spatial working memory, providing evidence against this possibility.

These two accounts make distinct predictions about responses in physical > social fROIs during a non-spatial working memory task (e.g. digit plan, mental arithmetic, verbal working memory). Prior research has reported greater similarity in patterns of activity within a pair of two demanding working memory tasks, and within a pair of two physical reasoning tasks, than across task sets (Fischer et al., 2016), but did not study whether activity in IP fROIs is modulated by task difficulty, as we report here. Under the hypothesis that the representations implemented in IP fROIs are useful for any demanding task, these fROIs should also respond more to difficult conditions of tasks from other domains. Under the hypothesis that representations of physical objects are also useful for tasks that place high demands on spatial computations, then these fROIs should respond equally to difficult and easy versions of non visuospatial tasks from other domains.

In general, the project of identifying and disentangling the domain-general and domain-specific components of physical reasoning (or any complex mental function) faces two interrelated conceptual challenges. The first challenge is identifying what representations and computations fall under each domain. For example, computational models of intuitive physics include variables like *body*, *mass*, and *velocity*, but also make use of variables like *distance*, *direction*, and *location* (Battaglia et al., 2013; Smith et al., 2019; Ullman et al., 2017). Are the first set of variables distinctive to intuitive physics, and the second set distinctive to spatial cognition? Do we use the same mental machinery to reason about the distance, direction and location of objects in a physical reasoning task, and to navigate from one location to another using a map? Proposing and revising the ontology of cognitive capacities is a critical step in the process of studying their neural mechanisms (Francken et al., 2022). A second related challenge is taking care to distinguish two claims: (i) the claim that mental capacity A is *irreducible* to mental capacity B, and (ii) the claim that mental capacity A is *independent* of mental capacity B. In the case of intuitive physics and spatial reasoning, we suspect that these two capacities are not redundant, but may very well be inter-related. Indeed, many physical reasoning tasks (e.g. judging the relative masses of two colliding objects; making a prediction about which way an unstable tower will tip; predicting where an object will move to in a dynamic scene) likely rely on both physical and spatial representations. Likewise, many spatial reasoning tasks (e.g. mental rotation, path integration, map reading), likely require use of physical representations of bodies, but perhaps not other variables like mass. Thus, finding that physical reasoning is not reducible to spatial reasoning should not be mistaken for evidence that these two domains are completely independent (Mitko & Fischer, 2020); likewise, the current findings should not be mistaken for evidence that there is no such thing as a domain-specific system for intuitive physics. The full empirical project of characterizing mental and neural architecture will require the field to study not only the boundaries between cognitive domains, but also how they interact.

### 4.2 Limitations and Conclusion

The current work has limitations. First, we were unable to study individual differences in the physical prediction task due to low reliability of the behavioral data. Therefore, it is an open question whether neural responses in MD and IP fROIs predict physical reasoning behaviors; future work could make use of reliable tasks of physical reasoning (Mitko et al., 2024) to address this issue. Second, it is unclear whether the lack of correspondence between the hard > easy responses in MD fROIs and spatial working memory behavior is spurious and/or underpowered. In prior research, Assem et al. (2020) found with a much larger sample size that individual differences in hard > easy spatial working memory responses in frontoparietal cortex correlated both with performance in the scanner, *and* with an independent test of fluid intelligence. Third, the nature of the individual differences in this paper is left open by our data. It is possible that the physical > social responses we studied also reflect a trait individual difference, a state individual difference, or a mix of both. Lastly, although we found that responses in physical > social fROIs predicted behavior in the spatial working memory task, a stronger test of the functional role of these regions in supporting behavior would come from studies using tools like transcranial magnetic stimulation (TMS). For example, it is possible that perturbing function in portions of the frontoparietal cortex maximally responsive during physical reasoning would only change performance in physical inference tasks; another possibility is that applying TMS over these regions would also modulate spatial reasoning, or performance in other demanding tasks.

In conclusion, our findings converge with the broader picture that areas of the frontal and parietal cortex support physical reasoning. We showed that the voxels in these areas whose activity is modulated by task difficulty are not strongly preferentially involved in physical (vs social) reasoning. Yet, the voxels in the same areas that are preferentially responsive during physical reasoning are just as responsive during a demanding spatial working memory task. This asymmetry in the function of these areas, and more broadly, the connection between physical reasoning and other cognitive domains, should continue to be important areas for future research.

## 5. Acknowledgements

Thank you to the members of the Liu Lab, Early Career Writing Group, Marina Bedny, and Leyla Isik for feedback on an earlier draft of this manuscript.

## Footnotes

1 We pre-registered this model with an additional random intercept for run as well, but this led to a singular fit, so we dropped this random intercept.

2 We opted to focus on accuracy, as opposed to reaction time, because the temporal window measuring button press responses differed substantially between tasks (3.75 seconds for the spatial working memory task, and 1.5 seconds for the physical prediction task).

