## Supplemental Materials for "Distinct and shared neural resources between processing of dynamic physical objects and spatial working memory"

This document contains supplemental figures and results associated with the manuscript "*Distinct and shared neural resources between processing of dynamic physical objects and spatial working memory*" by Samuel Maione and Shari Liu. Correspondence concerning this paper should be addressed to Samuel Maione.

**Supplemental Materials** **1**
1. Performance across two main tasks 2
2. Preprocessing of neural data (fMRIprep boilerplate) 3
3. Spatial profile of fROIs 5
   3.1 Dice's Coefficient: Additional tables and figures 5
   3.2 Spatial extent of suprathresholded voxels 7
   3.3 Lateralization indexes 8
4. Additional univariate analyses 10
   4.1 Univariate responses in fROIs removing spatial overlap 11
   4.2 Univariate responses in overlapping voxels between hard > easy and physical > social fROIs 13
   4.3 Univariate responses in suprathreshold (p < 0.001, uncorrected) hard > easy and physical > social voxels 15
5. Correlations between effects in MD and IP fROIs, and V1 16
6. Model selection for predictors of spatial working memory performance 18
7. Overlap causal mediation analysis 20
8. Robustness check:  Effect sizes across top N% fROI definitions 21

**1. Performance across two main tasks**

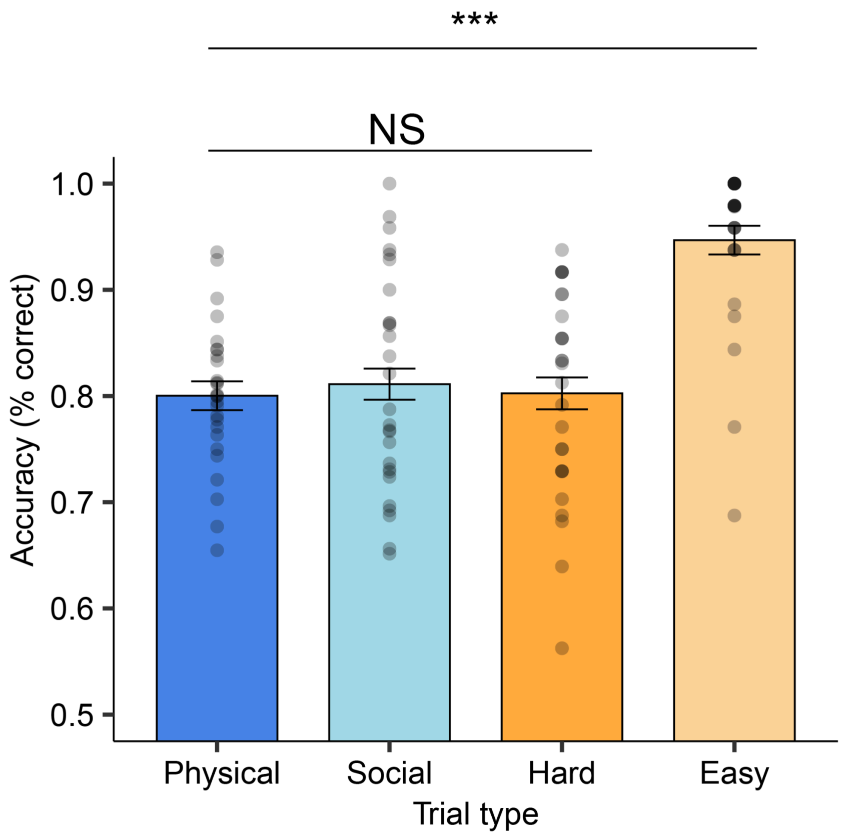


**Figure S1.** Performance on the physical vs social prediction task, and the hard vs easy spatial working memory task. Each data point corresponds to the average accuracy of a participant across two runs. Statistics derived from a regression modeling accuracy across conditions, `Performance ~ Trial_type + (1|Subject)`. NS indicates *p* >= .1; + < .1; * < .05; **< .01; *** < .001, two-tailed. Error bars indicate standard errors.

**2. Preprocessing of neural data (fMRIprep boilerplate)**

Results included in this manuscript come from preprocessing performed using **fMRIPrep 23.1.4**, which is based on **Nipype 1.8.6**.

**Anatomical data preprocessing**: A total of 1 T1-weighted (T1w) images were found within the input BIDS dataset.The T1-weighted image was corrected for intensity non-uniformity (INU) with `N4BiasFieldCorrection`, distributed with ANTs (version unknown), and used as T1w-reference throughout the workflow. The T1w-reference was then skull-stripped with a Nipype implementation of the `antsBrainExtraction.sh` workflow (from ANTs), using OASIS30ANTs as target template. Brain tissue segmentation of Cerebrospinal fluid (CSF), white-matter (WM) and gray-matter (GM) was performed on the brain-extracted T1w using `fast` (fsl). Brain surfaces were reconstructed using `recon-all` (FreeSurfer 7.3.2), and the brain mask estimated previously was refined with a custom variation of the method to reconcile ANTs-derived and FreeSurfer-derived segmentations of the cortical gray-matter of Mindboggle. Grayordinate “dscalar” files containing 91k samples were also generated using the highest-resolution `fsaverage` as an intermediate standardized surface space.

Volume-based spatial normalization to two standard spaces (MNI152NLin6Asym, MNI152NLin2009cAsym) was performed through nonlinear registration with `antsRegistration` (ANTs), using brain-extracted versions of both T1w reference and the T1w template. The following templates were were selected for spatial normalization and accessed with TemplateFlow (v23.0.0): FSL's MNI ICBM 152 non-linear 6th Generation Asymmetric Average Brain Stereotaxic Registration Model, ICBM 152 Nonlinear Asymmetrical template version 2009c.

**Functional data preprocessing**: First, a reference volume and its skull-stripped version were generated using a custom methodology of fMRIPrep. Head-motion parameters with respect to the BOLD reference (transformation matrices, and six corresponding rotation and translation parameters) are estimated before any spatiotemporal filtering using `mcflirt`. BOLD runs were slice-time corrected to 0.948s (0.5 of slice acquisition range 0s-1.9s) using `3dTshift` from AFNI. The BOLD time-series (including slice-timing correction when applied) were resampled onto their original, native space by applying the transforms to correct for head-motion. These resampled BOLD time-series will be referred to as preprocessed BOLD in original space, or just preprocessed BOLD. The BOLD reference was then co-registered to the T1w reference using `bbregister` (FreeSurfer) which implements boundary-based registration. Co-registration was configured with six degrees of freedom. Several confounding time-series were calculated based on the preprocessed BOLD: framewise displacement (FD), DVARS and three region-wise global signals. FD was computed using two formulations following Power (absolute sum of relative motions) and Jenkinson (relative root mean square displacement between affines). FD and DVARS are calculated for each functional run, both using their implementations in Nipype. The three global signals are extracted within the CSF, the WM, and the whole-brain masks. Additionally, a set of physiological regressors were extracted to allow for component-based noise correction. Principal components are estimated after high-pass filtering the preprocessed

BOLD time-series (using a discrete cosine filter with 128s cut-off) for the two CompCor variants: temporal (tCompCor) and anatomical (aCompCor). tCompCor components are then calculated from the top 2% variable voxels within the brain mask. For aCompCor, three probabilistic masks (CSF, WM and combined CSF+WM) are generated in anatomical space. The implementation differs from that of Behzadi et al. in that instead of eroding the masks by 2 pixels on BOLD space, a mask of pixels that likely contain a volume fraction of GM is subtracted from the aCompCor masks. This mask is obtained by dilating a GM mask extracted from the FreeSurfer's aseg segmentation, and it ensures components are not extracted from voxels containing a minimal fraction of GM. Finally, these masks are resampled into BOLD space and binarized by thresholding at 0.99 (as in the original implementation).

Components are also calculated separately within the WM and CSF masks. For each CompCor decomposition, the *k* components with the largest singular values are retained, such that the retained components' time series are sufficient to explain 50 percent of variance across the nuisance mask (CSF, WM, combined, or temporal). The remaining components are dropped from consideration. The head-motion estimates calculated in the correction step were also placed within the corresponding confounds file. The confound time series derived from head motion estimates and global signals were expanded with the inclusion of temporal derivatives and quadratic terms for each. Frames that exceeded a threshold of 0.5 mm FD or 1.5 standardized DVARS were annotated as motion outliers. Additional nuisance timeseries are calculated by means of principal components analysis of the signal found within a thin band (crown) of voxels around the edge of the brain, as proposed by Patriat et al. 2017. The BOLD time-series were resampled into standard space, generating a preprocessed BOLD run in MNI152NLin6Asym space. First, a reference volume and its skull-stripped version were generated using a custom methodology of fMRIPrep. The BOLD time-series were resampled onto the left/right-symmetric template "fsLR". Grayordinates files containing 91k samples were also generated using the highest-resolution `fsaverage` as intermediate standardized surface space. All resamplings can be performed with a single interpolation step by composing all the pertinent transformations (i.e. head-motion transform matrices, susceptibility distortion correction when available, and co-registrations to anatomical and output spaces). Gridded (volumetric) resamplings were performed using `antsApplyTransforms` (ANTs), configured with Lanczos interpolation to minimize the smoothing effects of other kernels. Non-gridded (surface) resamplings were performed using `mri_vol2surf` (FreeSurfer).

Many internal operations of fMRIPrep use Nilearn 0.10.1, mostly within the functional processing workflow. For more details of the pipeline, see https://fmriprep.readthedocs.io/en/latest/workflows.html.

The above boilerplate text was automatically generated by fMRIPrep with the express intention that users should copy and paste this text into their manuscripts unchanged. It is released under the CC0 license.

### 3. Spatial profile of fROIs

#### *3.1 Dice's Coefficient: Additional tables and figures*

**Table S1.** Adjusted Dice's coefficients by parcel, generated by dividing the empirical overlap between hard > easy and physical > social fROIs by the smallest DC we would expect, given overlap between two sets of hard > easy fROIs and physical > social fROIs, whichever is lower. See **section 3.2**.

| Parcel | Unscaled DC (Physical > social vs hard > easy) | Noise ceiling (source) | Scaled DC (Physical > social vs hard > easy) | DC *SE* |
|---|---|---|---|---|
| PPC_L | 0.157 | 0.430 (physical > social) | 0.364 | 0.043 |
| MPC_L | 0.208 | 0.485 (hard > easy) | 0.428 | 0.050 |
| APC_L | 0.161 | 0.324 (physical > social) | 0.497 | 0.044 |
| PC_L | 0.145 | 0.433 (physical > social) | 0.336 | 0.043 |
| PPC_R | 0.146 | 0.331 (physical > social) | 0.441 | 0.052 |
| MPC_R | 0.190 | 0.279 (physical > social) | 0.679 | 0.087 |
| APC_R | 0.203 | 0.269 (physical > social) | 0.754 | 0.069 |
| PC_R | 0.155 | 0.361 (physical > social) | 0.428 | 0.039 |

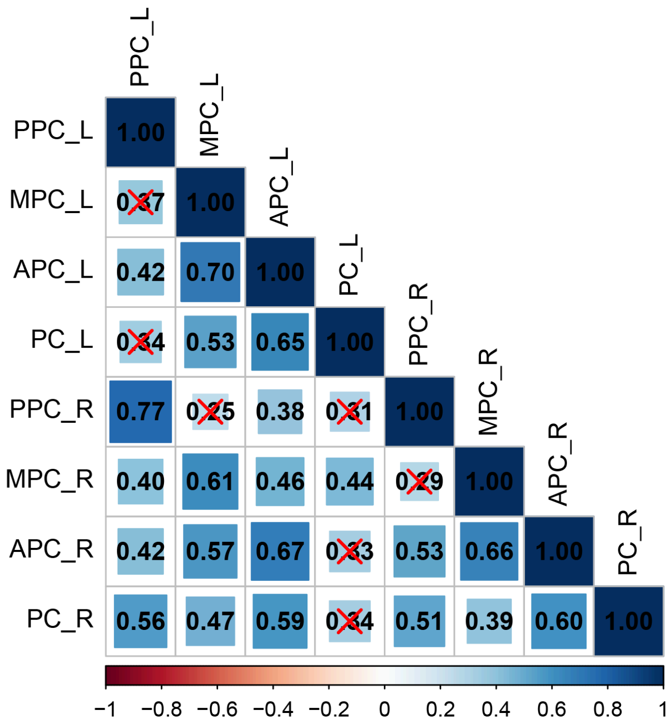


**Figure S2.** Correlation of Dice's coefficients (Spearman's rho) between fROIs (hard > easy vs. physical > social) split by parcel. Insignificant correlations marked with red 'X' ($p > 0.05$). Overall, the majority of DCs between fROIs within parcels correlated with one another, and thus we used the average DC for each participant for individual difference analyses.

#### *3.2 Spatial extent of suprathresholded voxels*

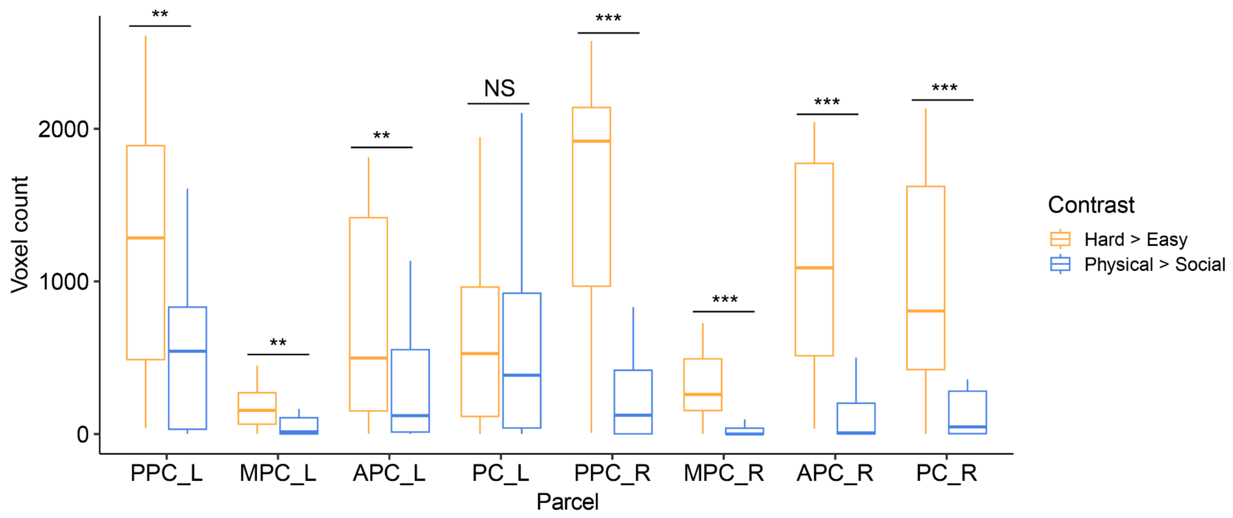


**Figure S3.** Number of hard > easy and physical > social suprathreshold voxels ($p < 0.001$; uncorrected) within each parcel in each person's data. Generally, we see more hard > easy voxels than physical > social voxels, suggesting that the hard > easy response has a greater spatial extent. (Note: Overlapping voxels are included in this plot.) Statistics computed with a linear model: `Voxel_count ~ contrast`.[1] NS indicates $p$ >= .1; + < .1; * < .05; **< .01; *** < .001, two-tailed.

[1] When we attempted to model the data including a random intercept for each participant, this led to a singular fit.

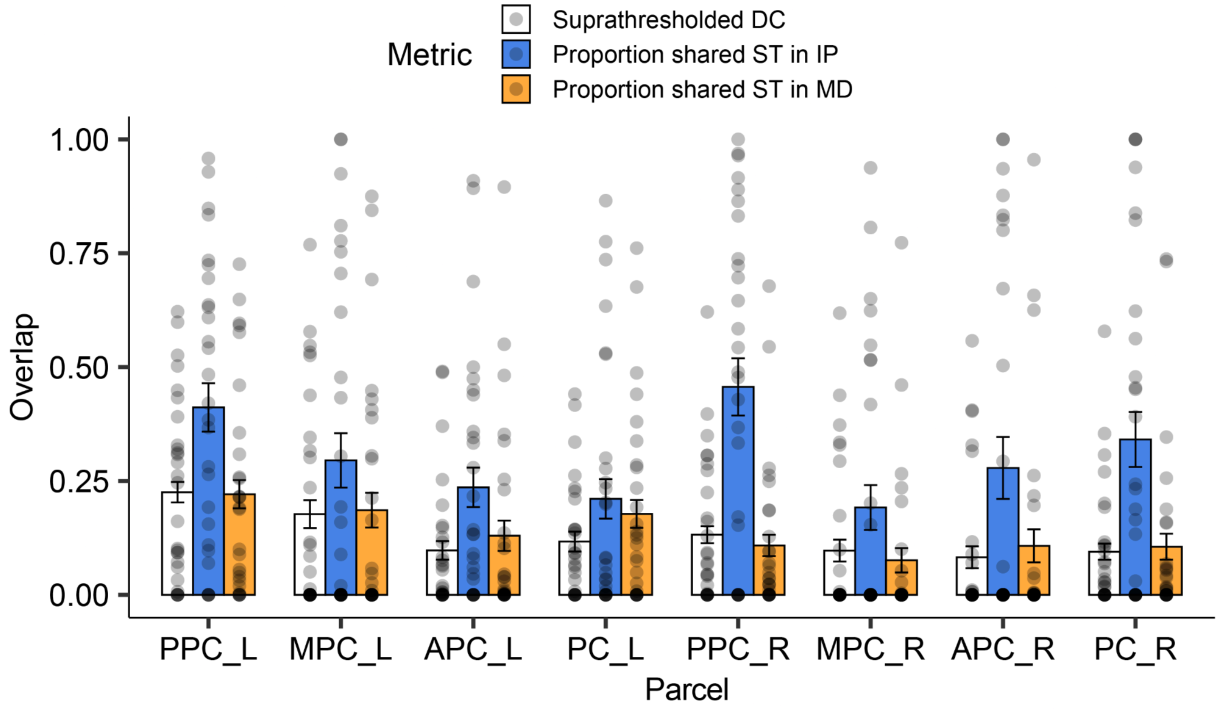


**Figure S4.** Three metrics characterizing spatial overlap in suprathreshold ($p < 0.001$, uncorrected) voxels within individual participants: Dice's coefficient (DC; white), the proportion of suprathreshold IP voxels that overlap with suprathreshold MD voxels (blue), and the proportion of suprathreshold MD voxels that overlap with IP voxels (orange).

***3.3 Lateralization indexes***

Using a conventional formula from (Brumer et al., 2020) the lateralization index (LI) is defined as:

$$LI = (LV - RV) / (LV + RV)$$

Where LV is the number of suprathreshold voxels in the left hemisphere and RV is the number of suprathreshold voxels in the right hemisphere. Like in the other analyses, we defined the threshold as $p < 0.001$, uncorrected.

For each subject, we computed LV and RV by constraining the search to the 4 anatomical ROIs (PPC, MPC, APC, PC in each hemisphere), and summing up the number of suprathreshold voxels across the frontoparietal cortex.

The IP regions were left lateralized ($B = 0.49$, 95% CI = [0.35, 0.63], $p < 0.001$, two-tailed), and MD regions were right lateralized ($B = -0.24$, 95% CI = [-0.10, -0.38], $p = 0.001$, two-tailed), linear model comparing distribution of LI to 0. These two LIs significantly differed ($B = 0.73$, 95%

CI = [0.53, 0.93], $p < 0.001$, two-tailed) as assessed by a linear model comparing distribution of LI across two contrasts[2].

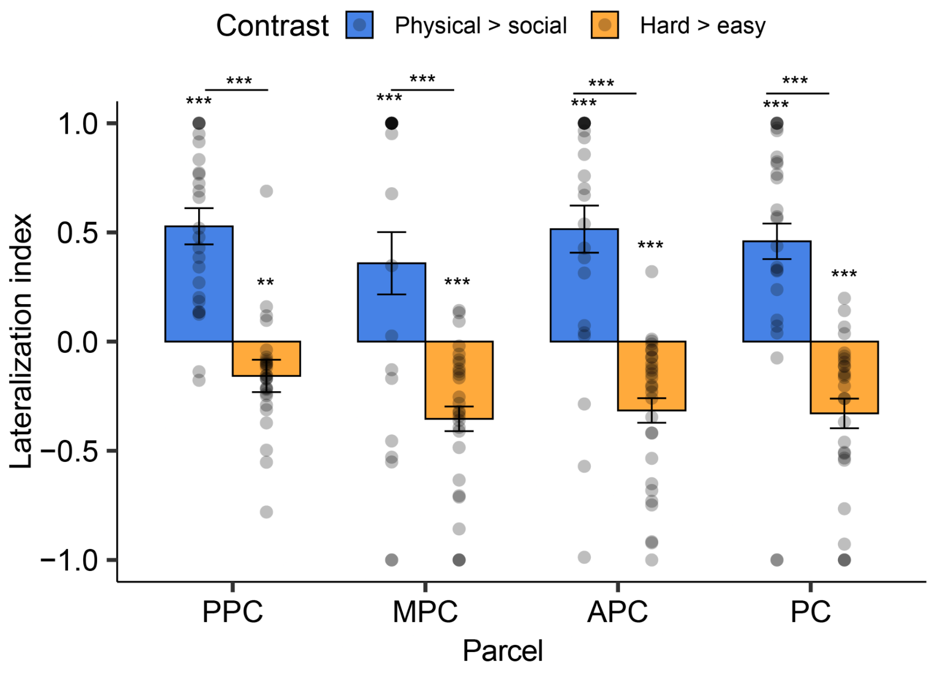


**Figure S5.** Lateralization index (LI) split by parcel (>0 indicates left lateralized; <0 indicates right lateralized). Comparisons above use the model formula `LI ~ contrast` to assess significance between contrasts and significance from zero. NS indicates *p* >= .1; + < .1; * < .05; **< .01; *** < .001, two-tailed. Error bars indicate standard errors.

[2] When we attempted to model the data including a random intercept for each participant, this led to a singular fit.

**4. Additional univariate analyses**

**Table S2.** Comparing responses of MD fROIs and IP fROIs within the same tasks. Significant *p*-values (threshold of $\alpha$ = .05, two-tailed) are bolded. * indicates results from individual regions that pass a more conservative corrected threshold of $\alpha$ = .00625 (8 fROIs per category).

| | | hard > easy (MD fROIs > IP fROIs) | | physical > social (IP fROIs > MD fROIs) | |
|---|---|---|---|---|---|
| Parcel | df | Std. Beta | *p*-value | Std. Beta | *p*-value |
| PPC_L | 193 | 0.24 | 0.136 | 0.26 | 0.077 |
| MPC_L | 193 | 0.20 | 0.266 | 0.16 | 0.289 |
| APC_L | 193 | 0.44 | **0.008** | 0.18 | 0.214 |
| PC_L | 193 | 0.44 | **0.006*** | 0.36 | **0.017** |
| PPC_R | 193 | 0.36 | **0.022** | 0.13 | 0.392 |
| MPC_R | 193 | 0.32 | 0.066 | 0.04 | 0.826 |
| APC_R | 193 | 0.46 | **0.012** | 0.08 | 0.632 |
| PC_R | 193 | 0.55 | **0.001*** | 0.27 | 0.101 |
| all parcels | 1754 | 0.30 | **<0.001** | 0.15 | **0.020** |

In the main text, we compared the neural responses of the same fROIs to two different tasks. Here, we took an adjacent approach: in each parcel, we compared the magnitude of neural responses (hard > easy or physical > social) in the canonical set of fROIs vs the opposite set of fROIs. Put another way, this table measures the response-by-fROI set interaction, such that an interaction denotes a neural response is larger in one set of regions than the other.

#### *4.1 Univariate responses in fROIs removing spatial overlap*

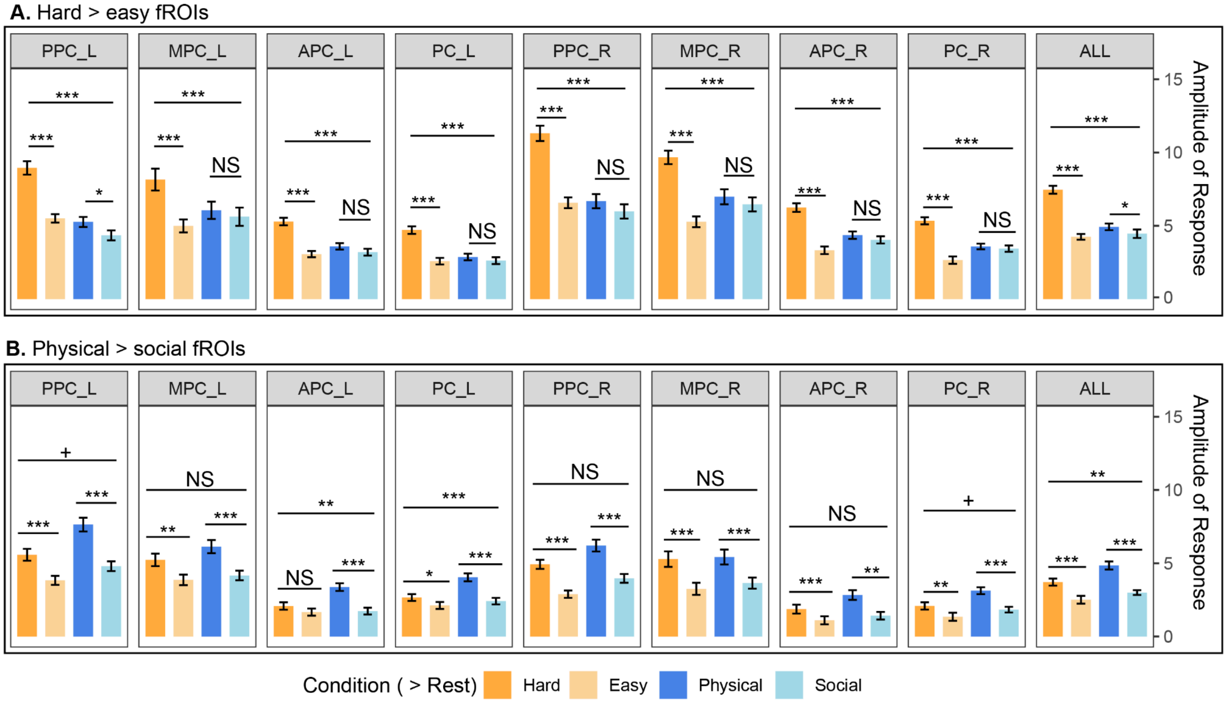


**Figure S6.** Responses in physical > social and hard > easy fROIs (top 10%), removing voxels that overlapped between fROIs within a parcel. We used the same model formula from the corresponding main-text analysis: `beta ~ preference * task + (1|subject)`. See **Section 2.7.3** for details. Overall, we see the same results as reported in the main text: by our strictest significance threshold, a greater hard > easy than physical > social response in all 8 hard > easy (MD) fROIs, and a greater physical > social than hard > easy response in only 2 of 8 physical > social (IP) fROIs. These two IP fROIs, left anterior parietal cortex (APC_L) and left precentral cortex (PC_L), showed a significant task-by-preference interaction in this analysis, but no such interaction in the main analysis. Error bars indicate +/-1 standard error, NS indicates p >= .1; + < .1; * < .05; **< .01; *** < .001, two-tailed.

**Table S3.** Univariate responses to both tasks in hard > easy (MD) and physical > social (IP) fROIs removing overlapping voxels. Significant p-values (threshold of $\alpha$ = .05, two-tailed) are bolded. * indicates significant p-values from individual regions that pass a more conservative corrected threshold of $\alpha$ = .00625 (8 fROIs per ROI set, MD and IP).

| | | Physical prediction > social prediction | | Hard > easy spatial working memory | | Interaction | |
|---|---|---|---|---|---|---|---|
| fROI | df | Std. Beta | *p*-value | Std. Beta | *p*-value | Std. Beta | *p*-value |
| MD PPC_L | 193 | 0.23 | **0.021** | 0.87 | **<0.001*** | 0.64 | **<0.001*** |
| MD MPC_L | 193 | 0.08 | 0.326 | 0.55 | **<0.001*** | 0.48 | **<0.001*** |
| MD APC_L | 193 | 0.16 | 0.141 | 0.87 | **<0.001*** | 0.72 | **<0.001*** |
| MD PC_L | 193 | 0.12 | 0.298 | 0.98 | **<0.001*** | 0.87 | **<0.001*** |
| MD PPC_R | 193 | 0.14 | 0.167 | 0.97 | **<0.001*** | 0.83 | **<0.001*** |
| MD MPC_R | 193 | 0.11 | 0.265 | 0.95 | **<0.001*** | 0.84 | **<0.001*** |
| MD APC_R | 193 | 0.12 | 0.302 | 1.10 | **<0.001*** | 0.98 | **<0.001*** |
| MD PC_R | 193 | 0.07 | 0.547 | 1.29 | **<0.001*** | 1.22 | **<0.001*** |
| MD all fROIs | 1754 | 0.11 | **0.012** | 0.78 | **<0.001*** | 0.67 | **<0.001*** |
| IP PPC_L | 193 | 0.70 | **<0.001*** | 0.43 | **<0.001*** | 0.27 | 0.053 |
| IP MPC_L | 193 | 0.50 | **<0.001*** | 0.35 | **0.001*** | 0.15 | 0.299 |
| IP APC_L | 193 | 0.70 | **<0.001*** | 0.18 | 0.137 | 0.52 | **0.002*** |
| IP PC_L | 193 | 0.77 | **<0.001*** | 0.25 | **0.018** | 0.51 | **0.001*** |
| IP PPC_R | 193 | 0.63 | **<0.001*** | 0.58 | **<0.001*** | 0.06 | 0.713 |
| IP MPC_R | 193 | 0.45 | **0.002*** | 0.51 | **<0.001*** | -0.06 | 0.761 |
| IP APC_R | 193 | 0.63 | **<0.001*** | 0.34 | **0.007** | 0.29 | 0.104 |
| IP PC_R | 193 | 0.69 | **<0.001*** | 0.40 | **0.001*** | 0.30 | 0.068 |
| IP all fROIs | 1754 | 0.54 | **<0.001** | 0.35 | **<0.001** | 0.19 | **0.004** |

#### *4.2 Univariate responses in overlapping voxels between hard > easy and physical > social fROIs*

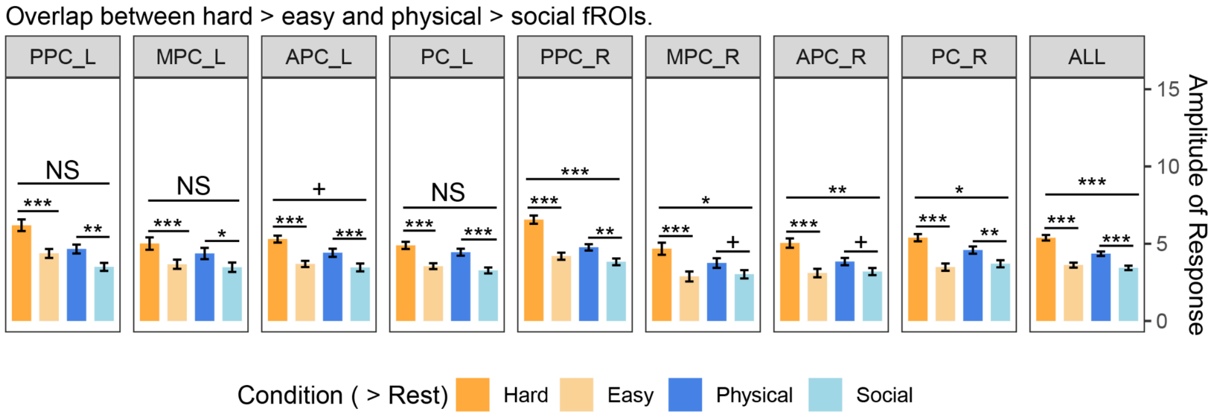


**Figure S7.** Responses including only overlapping voxels between physical > social and hard > easy fROIs. We used the same model formula from the corresponding main-text analysis: `beta ~ preference * task + (1|subject)`. See **Section 2.7.3** for details. Overall, we see that overlapping voxels in the right hemisphere showed a greater hard > easy than physical > social response, but overlapping voxels in the left hemisphere showed equally strong hard > easy and physical > social responses. Error bars indicate +/-1 standard error, NS indicates p >= .1; + < .1; * < .05; **< .01; *** < .001, two-tailed.

**Table S4.** Univariate responses to both tasks in voxels that overlap between hard > easy and physical > social fROIs. Significant *p*-values (threshold of $\alpha$ = .05, two-tailed) are bolded. * indicates significant p-values from individual regions that pass a more conservative corrected threshold of $\alpha$ = .00625 (8 regions).

| | | Physical prediction > social prediction | | Hard > easy spatial working memory | | Interaction | |
|---|---|---|---|---|---|---|---|
| fROI | df | Std. $\beta$ | *p*-value | Std. $\beta$ | *p*-value | Std. $\beta$ | *p*-value |
| PPC_L | 193 | 0.41 | **0.001*** | 0.64 | **<0.001*** | 0.23 | 0.190 |
| MPC_L | 193 | 0.26 | **0.013** | 0.39 | **<0.001*** | 0.13 | 0.370 |
| APC_L | 193 | 0.38 | **0.001*** | 0.64 | **<0.001*** | 0.26 | 0.093 |
| PC_L | 193 | 0.48 | **<0.001*** | 0.55 | **<0.001*** | 0.07 | 0.658 |
| PPC_R | 193 | 0.36 | **0.002*** | 0.92 | **<0.001*** | 0.55 | **0.001*** |
| MPC_R | 193 | 0.25 | 0.054 | 0.62 | **<0.001*** | 0.37 | **0.044** |
| APC_R | 193 | 0.27 | 0.058 | 0.82 | **<0.001*** | 0.55 | **0.007** |
| PC_R | 193 | 0.38 | **0.003*** | 0.83 | **<0.001*** | 0.45 | **0.016** |
| all fROIs | 1754 | 0.34 | **<0.001** | 0.65 | **<0.001** | 0.31 | **<0.001** |

#### *4.3 Univariate responses in suprathreshold (p < 0.001, uncorrected) hard > easy and physical > social voxels*

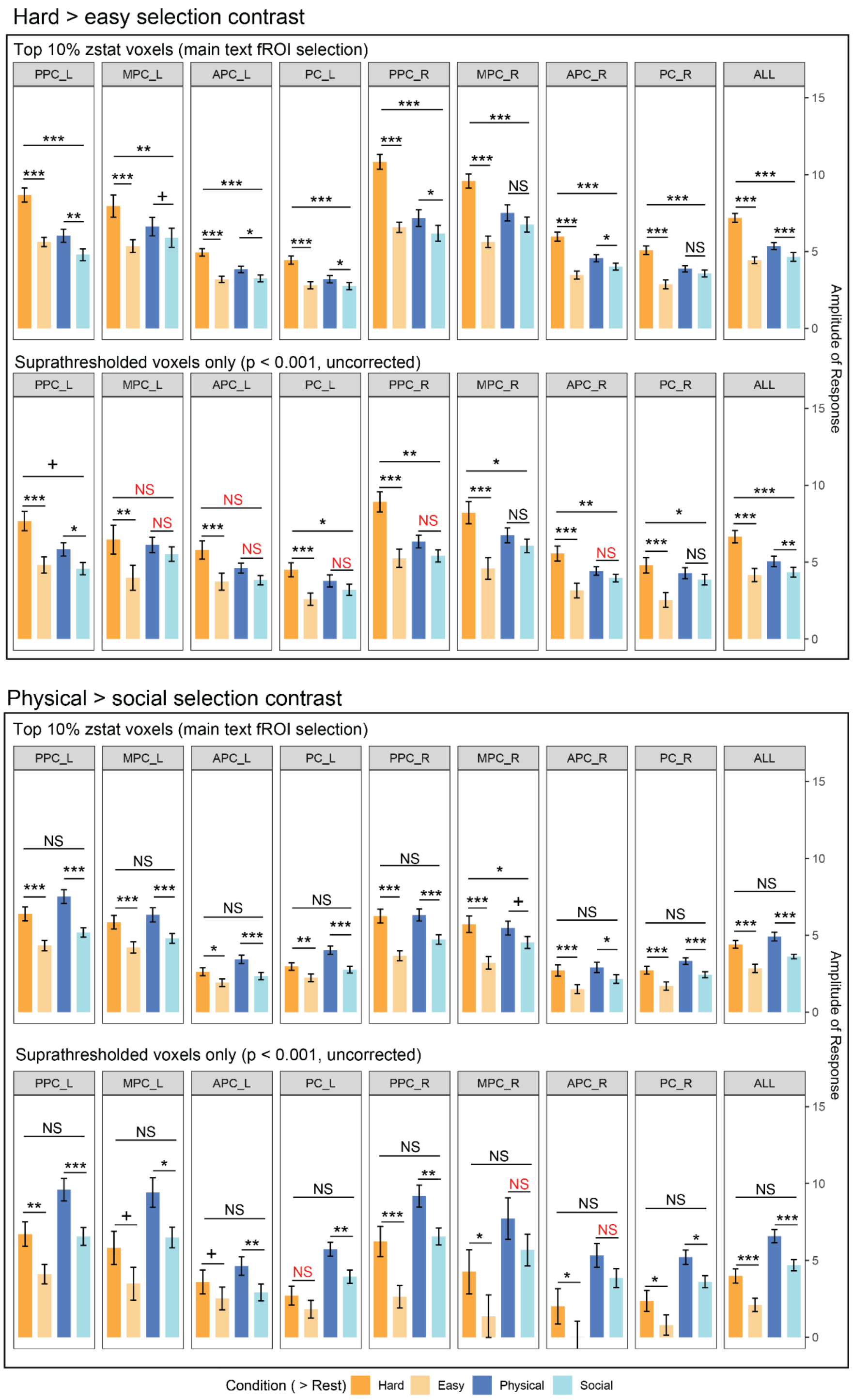


**Figure S8.** Comparison of univariate results in top 10% fROIs, as in the main text (both panels, top row) or suprathreshold voxels (both panels, bottom row). Statistics that flip significance (e.g. insignificant to significant or vice-versa) are colored in red. We used the same model formula from the corresponding main-text analysis: `beta ~ preference * task + (1|subject)`. See **Section 2.7.3** for details. Note that analyses in suprathreshold voxels include fewer observations, and a different number of observations across participants and regions. NS indicates *p* >= .1; + < .1; * < .05; **< .01; *** < .001, two-tailed. Error bars indicate standard errors.

Overall we see similar, though weaker, results in suprathreshold voxels: a task-by-condition interaction in MD regions (top panel), and no such interaction in IP regions (bottom panel).

**5. Correlations between effects in MD and IP fROIs, and V1**

**Table S5.** Correlations in hard > easy and physical > social responses between regions in the frontoparietal cortex and stimulus > rest responses in V1. Model formula: `correlation ~ pair_type`, where pair_type is either IP-MD (reference level, correlations across fROI sets), or FP-V1 (including both correlations between IP and MD fROIs to V1). The main finding is that the correlation between responses in IP and MD fROIs is significantly greater than the correspondence between either fROI set to V1.

| Task | Network Pair (FP indicates both MD and IP) | *df* | *t* | Unstandardized *B* coefficient [95% CI] | *p*-value |
|---|---|---|---|---|---|
| Model 1: Physical prediction (physical > social) | IP-MD (intercept) | 70 | 34.54 | 0.53 [0.50, 0.56] | **<0.001** |
| | FP-V1 vs. IP-MD | 70 | -20.75 | -0.67 [-0.74, -0.61] | **<0.001** |
| Model 2: Spatial working memory (hard > easy) | IP-MD (intercept) | 70 | 30.60 | 0.55 [0.52, 0.59] | **<0.001** |
| | FP-V1 vs. IP-MD | 70 | -10.79 | -0.41 [-0.49, -0.34] | **<0.001** |
| Model 3: Cross-task; physical > social in IP regions, hard > easy in MD regions | IP-MD (intercept) | 70 | 14.90 | 0.27 [0.24, 0.31] | **<0.001** |
| | FP-V1 vs. IP-MD | 70 | -4.31 | -0.17 [-0.25, -0.09] | **<0.001** |

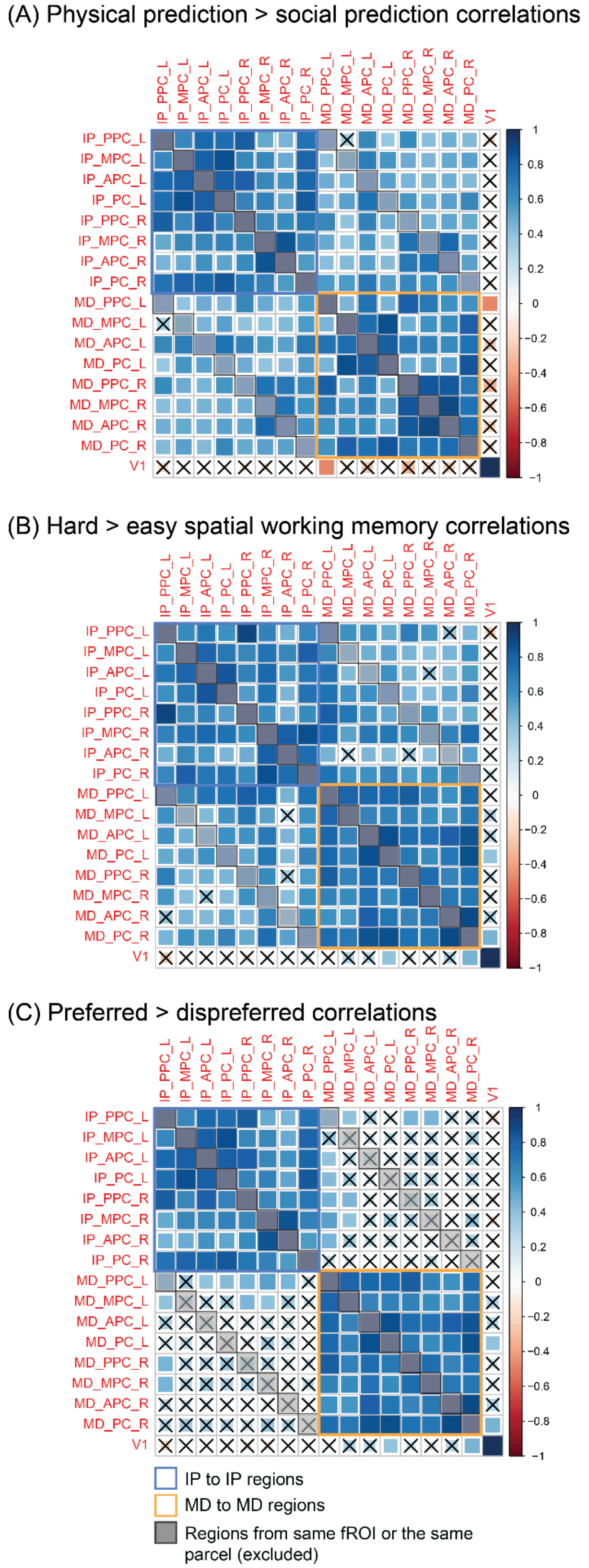


**Figure S9**. Correlation matrix (Spearman's rho) comparing hard > easy responses (A) and physical > social responses (B), and between these two responses (C) within and between fROI sets. Grey boxes indicate fROIs within a parcel, which were excluded from analyses on these data; 'X' denotes insignificant correlation between a pair of regions ($p > 0.05$, two-tailed). These effects are summarized in the main text, **Figure 5**, and reported in **Section 3.4**.

### 6. Model selection for predictors of spatial working memory performance

**Table S6.** Stepwise model results using all possible predictors. The best model, highlighted in green, is reported in the main text.

| Model | *AIC* | *BIC* | *df* | *F* | *p* |
|---|---|---|---|---|---|
| 1 | -45.96 | -43.37 | 26 | n/a | n/a |
| 2 | -47.56 | -47.32 | 25 | 4.64 | **0.042** |
| 3 | -52.50 | -47.32 | 24 | 7.38 | **0.012** |
| 4 | -52.53 | -46.06 | 23 | 0.91 | 0.352 |
| 5 | -50.54 | -42.76 | 22 | 0.92 | 0.348 |
| 6 | -51.40 | -42.33 | 21 | 2.35 | 0.140 |

Model 1: `hard_accuracy ~ 1`
Model 2: `hard_accuracy ~ MD_hard>easy`
Model 3: `hard_accuracy ~ MD_hard>easy + IP_hard>easy`
Model 4: `hard_accuracy ~ MD_hard>easy + IP_hard>easy + MD_phys>soc`
Model 5: `hard_accuracy ~ MD_hard>easy + IP_hard>easy + MD_phys>soc + IP_phys>soc`
Model 5: `hard_accuracy ~ MD_hard>easy + IP_hard>easy + MD_phys>soc + IP_phys>soc + V1_stim>rest`

**Table S7.** Stepwise model results using significant zero order correlations (see **Figure 6**) as predictors. The best model, highlighted in green, is reported in the main text.

| Model | *AIC* | *BIC* | *df* | *F* | *p* |
|---|---|---|---|---|---|
| 1 | -45.96 | -43.37 | 26 | n/a | n/a |
| 2 | -53.50 | -49.61 | 25 | 10.81 | **0.003** |
| 3 | -53.38 | -48.20 | 24 | 1.72 | 0.203 |
| 4 | -52.31 | -45.83 | 23 | 0.80 | 0.379 |

Model 1: `hard_accuracy ~ 1`
Model 2: `hard_accuracy ~ IP_hard>easy`
Model 3: `hard_accuracy ~ IP_hard>easy + MD_phys>soc`
Model 4: `hard_accuracy ~ IP_hard>easy + MD_phys>soc + IP_phys>soc`

**Table S8.** Stepwise model results comparing IP hard>easy alone to both IP hard>easy and MD hard>easy. We find IP hard>easy (alone) is the best model.

| Model | *AIC* | *BIC* | *df* | *F* | *p* |
|---|---|---|---|---|---|
| 1 | -45.96 | -43.37 | 26 | n/a | n/a |
| 2 | -53.50 | -49.61 | 25 | 10.55 | **0.003** |
| 3 | -52.50 | -47.32 | 24 | 0.91 | 0.350 |

Model 1: `hard_accuracy ~ 1`

Model 2: `hard_accuracy ~ IP_hard>easy`

Model 3: `hard_accuracy ~ IP_hard>easy + MD_hard>easy`

**7. Overlap causal mediation analysis**

Using the mediation package in R, we found the relationship between IP hard>easy responses and performance on the hard blocks of the spatial working memory task was mediated by the overlap across fROIs. In an exploratory analysis, we used 5000 simulations (Monte Carlo draws, nonparametric bootstrapping) to model the mediating effect of Dice's coefficient (DC). Across simulations, there was a significant average causal mediation effect (ACME) of spWM accuracy explained by fROI DC ($\beta$ = 0.22,95% CI = [0.01, 0.46], $p$ = 0.046, two-tailed). About 40.0% (95% CI = [8.7, 111.2], $p$ = 0.048) of the relationship between hard > easy responses in IP fROIs and behavioral performance was mediated by the degree of overlap between IP and MD regions.

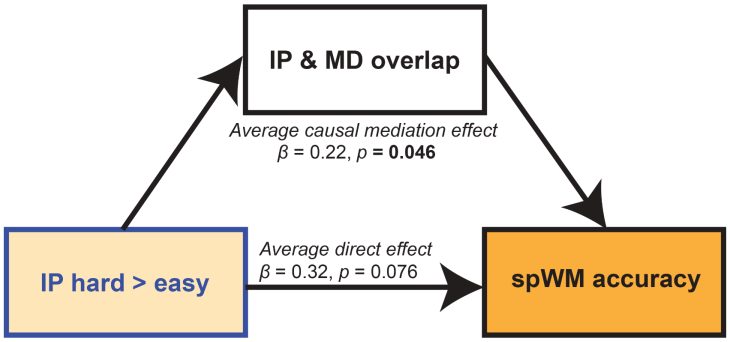


**Figure S10.** Proposed causal mediation relationship between IP hard > easy signal, overlap between IP and MD regions, and accuracy on the hard condition of the spatial working memory task.

**8. Robustness check: Effect sizes across top N% fROI definitions**

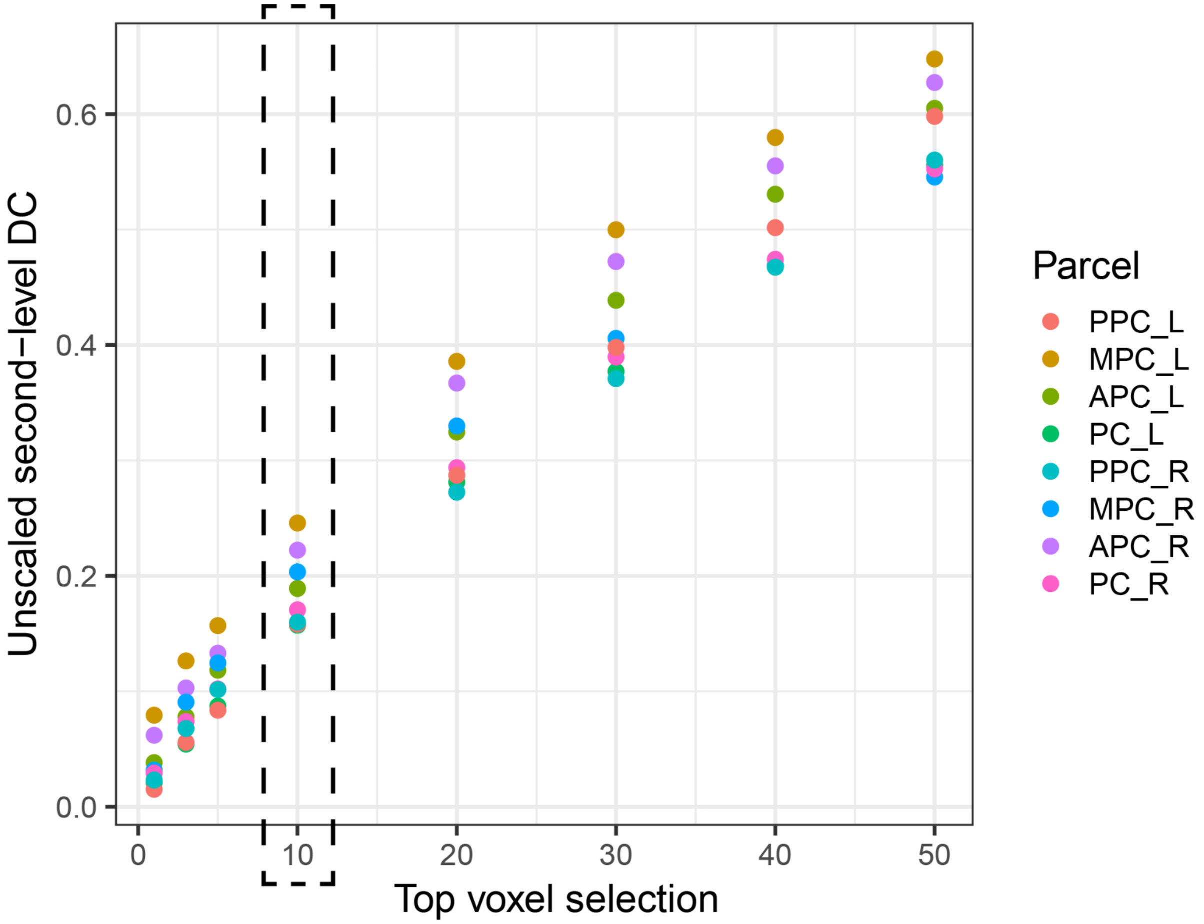


**Figure S11.** Dice's Coefficient between hard > easy and physical > social fROIs across different fROI selection methods (top 1, 3, 5, 10, 20, 30, 40, and 50%). Overall, this figure shows that DC increases approximately linearly with fROI size.

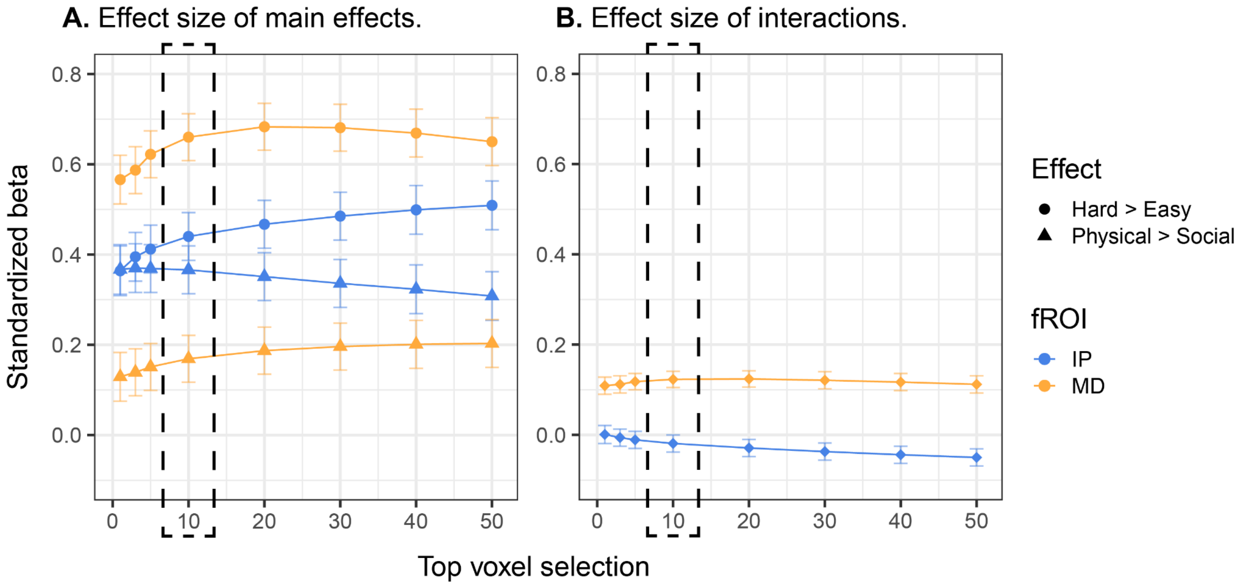


**Figure S12.** Effect size in standardized betas of (A) hard > easy responses, physics > social responses, and (B) their interaction, across varying top N% fROI definitions (top 10% was the pre-registered fROI size). Error bars denote +/- 1 standard error. Overall, we see that the effects remain relatively stable across fROI size.

At each fROI size (top 1, 3, 5, 10, 20, 30, 40, and 50%) we used a linear mixed effects model to compute the effect sizes of our three effects (`beta ~ condition * fROI_set + (1|ROI) + (1|subject)`) that were then standardized using `standardize_parameters()`. We looked at two main effects: (1) the effect size of a set of regions preference for hard trials relative to easy trials; and (2) the effect size of a set of regions preference for physical trials relative to social trials; and (3) interaction effect between the two main effects. Overall our results vary smoothly with fROI size.

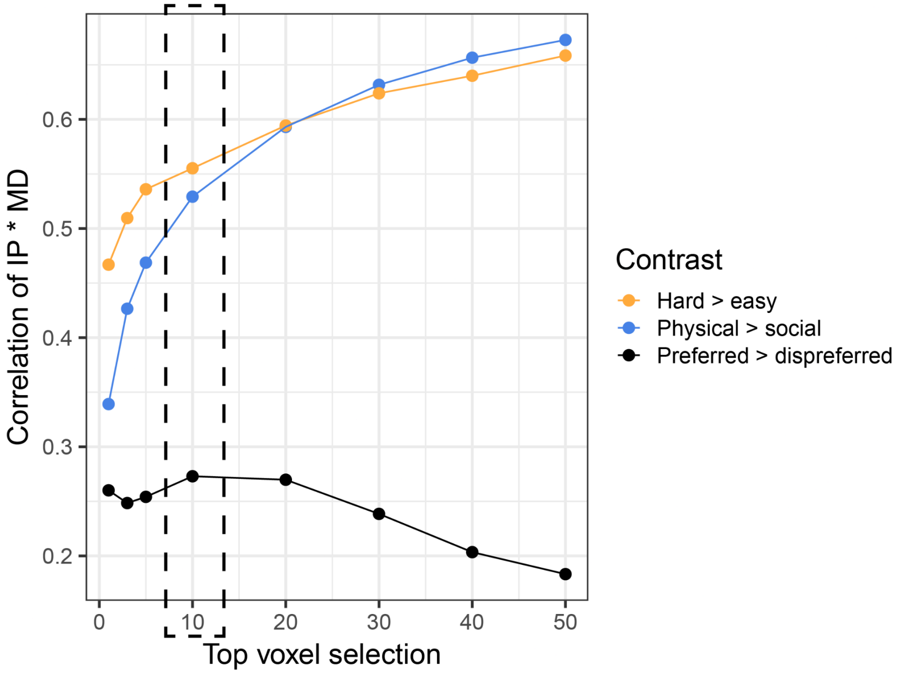


**Figure S13.** Spearman's ρ between univariate responses across fROI sets (corresponds to middle columns in **Figure 5A-C**).

At each fROI size (top 1, 3, 5, 10, 20, 30, 40, and 50%) we reported the correlation across the two sets of fROIs using hard > easy responses (orange), physical > social responses (blue), and preferred > dispreferred responses (i.e. cross-task; black). Overall our results vary smoothly with fROI size.

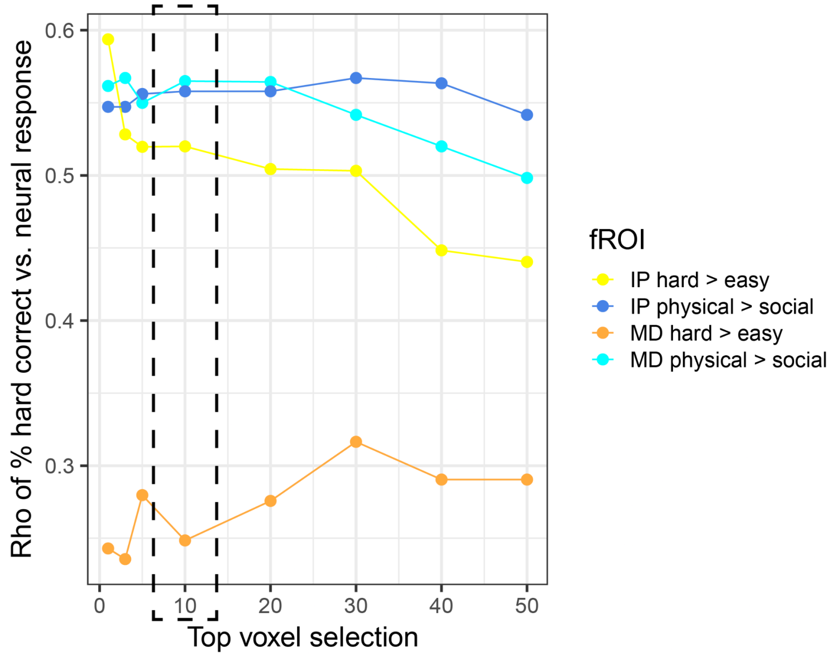


**Figure S14.** Spearman's ρ between performance on spatial working memory task and neural signal across fROI selection size.

At each fROI size (top 1, 3, 5, 10, 20, 30, 40, and 50%) we reported the correlation between accuracy on the spatial working memory task and responses from IP and MD fROIs. Overall our results vary smoothly with fROI size.

**Work cited**

Brumer, I., De Vita, E., Ashmore, J., Jarosz, J., & Borri, M. (2020). Implementation of clinically relevant and robust fMRI-based language lateralization: Choosing the laterality index calculation method. *PloS One*, *15*(3), e0230129.
